# Population dynamics underlying associative learning in the dorsal and ventral hippocampus

**DOI:** 10.1101/2021.11.16.468862

**Authors:** Jeremy S. Biane, Max A. Ladow, Fabio Stefanini, Sayi P. Boddu, Austin Fan, Shazreh Hassan, Naz Dundar, Daniel L. Apodaca-Montano, Nicholas I. Woods, Mazen A. Kheirbek

## Abstract

Animals associate cues with outcomes and continually update these associations as new information is presented. The hippocampus is crucial for this, yet how neurons track changes in cue-outcome associations remains unclear. Using 2-photon calcium imaging, we tracked the same dCA1 and vCA1 neurons across days to determine how responses evolve across phases of odor-outcome learning. We find that, initially, odors elicited robust responses in dCA1, whereas in vCA1 responses emerged after learning, including broad representations that stretched across cue, trace, and outcome periods. Population dynamics in both regions rapidly reorganized with learning, then stabilized into ensembles that stored odor representations for days, even after extinction or pairing with a different outcome. Finally, we found stable, robust signals across CA1 when anticipating reward, but not when anticipating inescapable shock. These results identify how the hippocampus encodes, stores, and updates learned associations, and illuminates the unique contributions of dorsal and ventral hippocampus.

## INTRODUCTION

As a child, an unexpected encounter with an ice cream truck can be a highly rewarding experience. To better predict the circumstances that led to this occurrence, the brain gathers information surrounding the incident, from broad cues associated with the availability of reward (the presence of music, the neighborhood in which the encounter occurred), to more detailed stimulus representations (the specific melody played, the precise location of the encounter), to the positive outcome from the experience of consuming ice cream. Following repeated encounters, the most predictive features are identified and used to inform behavior, such as running outside when a particular melody is heard.

This example illustrates a fundamental objective of the brain: to extract the underlying structure of the world and model its causal relationships. Moreover, the brain must be able to flexibly update these models as cue-outcome relationships change (e.g., when the melody is replaced, or the truck no longer carries your favorite ice cream). While the importance of examining the population dynamics underlying cognitive processes is becoming increasingly appreciated (Ahmed et al., 2020; Ebitz and Hayden, 2021; Stefanini et al., 2020), it is still generally unclear how learned associations are represented at the population level and how these representations change as a function of learning.

One area heavily implicated in encoding learned associations is the hippocampus. Genetic, anatomical, and functional data suggest the dorsal and ventral subdivisions of the rodent HPC (dHPC and vHPC) play distinct roles when learning about the world (Fanselow and Dong, 2010; Strange et al., 2014). Previous studies show neuronal responses in dHPC are relatively specific, encoding properties such as position within an environment, elapsed time, the identity of individual stimuli, and conjunctive representations such as object-location couplings (Kay and Frank, 2019; Komorowski et al., 2009; O’Keefe and Dostrovsky, 1971; Pastalkova et al., 2008; Taxidis et al., 2020; Wood et al., 2000; Yu et al., 2018). In contrast, vHPC representations respond to more abstract elements that generalize across distinct objects and events (Harland et al., 2017; Knudsen and Wallis, 2021; Komorowski et al., 2013; Royer et al., 2010) and tend to reflect the overall valence of an experience (Ciocchi et al.,2015; Jimenez et al., 2018).

While dHPC and vHPC may encode unique features of an explored environment, it remains unknown how these areas may be differentially engaged during the encoding of associative memories. Distinct encoding properties in dHPC and vHPC could not only enrich the scope of internal models, but also facilitate learning (Collin et al., 2015; Harland et al., 2017). For instance, during goal-oriented navigation, models incorporating both small-and large-scale place fields (characteristic of dHPC and vHPC, respectively) lead to faster learning compared to models only incorporating a single scale (Llofriu et al., 2015). During learning, detailed representations by dHPC may support the formation of associative memories on the basis of local cues, such as the precise identity of an object in an environment, while broad vHPC representations may generalize knowledge across multiple experiences and/or attach significance to contexts in which associations occur.

Here, we used 2-photon in vivo imaging of population activity in dCA1 or vCA1 to track the activity of the same neurons across multiple stages of learning as mice learned to associate odor stimuli with appetitive or aversive outcomes. This allowed us to examine how task-related information is differentially represented across the dorsoventral hippocampal axis and how these representations evolve with learning. Further, we examined the stability of representations across training, the influence of different outcomes on these encoding properties, and how neural representations adapt when cue-outcome relationships are altered.

## RESULTS

### Representations of odor identity across the dorsoventral axis of CA1

We imaged odor-evoked neural responses in dCA1 and vCA1 using high resolution 2-photon microscopy in mice expressing the calcium indicator GCaMP6f (Figure 1A-1C). In dCA1, we found that odors elicited a robust response in a subset of neurons and that odor-evoked population responses could be discriminated from ITI (baseline) activity with high accuracy using linear decoders (Figures 1E-1H, S1E). Moreover, odor identity could be accurately decoded from dCA1 population activity (Figure 1I, 1J). Surprisingly, however, odors did not elicit robust responses in vCA1 neurons, and linear decoders performed significantly worse compared to dCA1 when discriminating odor activity from baseline activity, or when reading out odor identity (Figures 1E-1J, S1D). This suggests that, during initial exposure to odorants, odor identities are reliably represented in dCA1 but not in vCA1.

**Figure 1.**
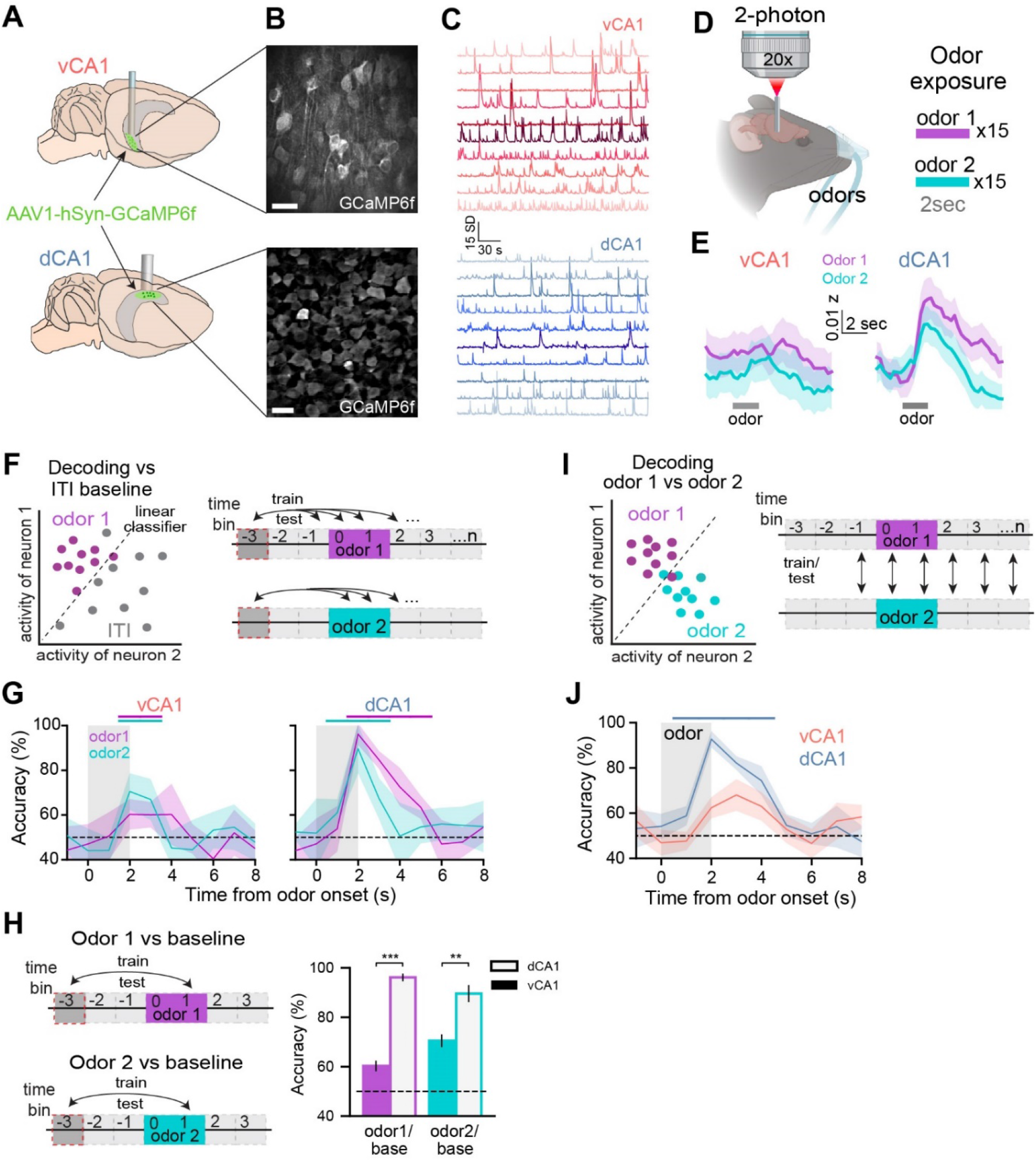
Prior to conditioning, odor stimuli are strongly represented in dCA1 but not vCA1. (A) AAV virus expressing GCaMP6f was targeted to vCA1 or dCA1, and a GRIN lens was implanted above the injection site. (B and C) Sample FOVs demonstrating (B) GCaMP expression and (C) time series data of denoised fluorescent traces. Scale bar in (B) = 25 µm. (D) Calcium signals were imaged while mice received 30 trials of 2 sec odor exposures (15 of each odor). (E) Population mean (±SEM) of z-scored fluorescent signals occurring around the onset of odor1 (purple) or odor2 (cyan). Grey bar = odor delivery period. (F) (left) Schematic of decoding procedure. Each dot represents the single-trial population activity vector during baseline (grey) or odor delivery (purple) periods. (right) Linear classifiers were trained to classify population activity patterns occurring during baseline from those occurring at time bin t. (G) Population-activity decoding accuracy for odor 1 or odor 2 from baseline (±SD). Colored-coded bars above graph denote periods where accuracy is significantly greater than chance (p < 0.01, Mann-Whitney U test). (H) Odor-period decoding. Pop. activity during the last second of odor delivery was used to decode odor 1 or odor 2 from baseline. (I and J) Same as (F) and (G) above, but decoding for trial type (odor 1 or odor 2) at each time bin t. For all figures: * p< 0.05, ** p < 0.01, *** p < 0.001. See Table S1 for all statistical analysis details.

### Attaching behavioral significance to odors enhances their representations in vCA1

Given the role of the ventral hippocampus in emotional and motivational processes (Tannenholz et al., 2014), we reasoned that odor representations may be enhanced in vCA1 if paired with a salient outcome. To disambiguate odor representations from potential reward anticipation signals, we used a two-odor trace appetitive conditioning paradigm wherein the CS+ odor was separated from sucrose reward delivery by a 2s trace period. Following ∼4 days of training, mice displayed anticipatory licks during the CS+ trace period, with minimal licking during all other task periods (Figure 2A-2D).

**Figure 2.**
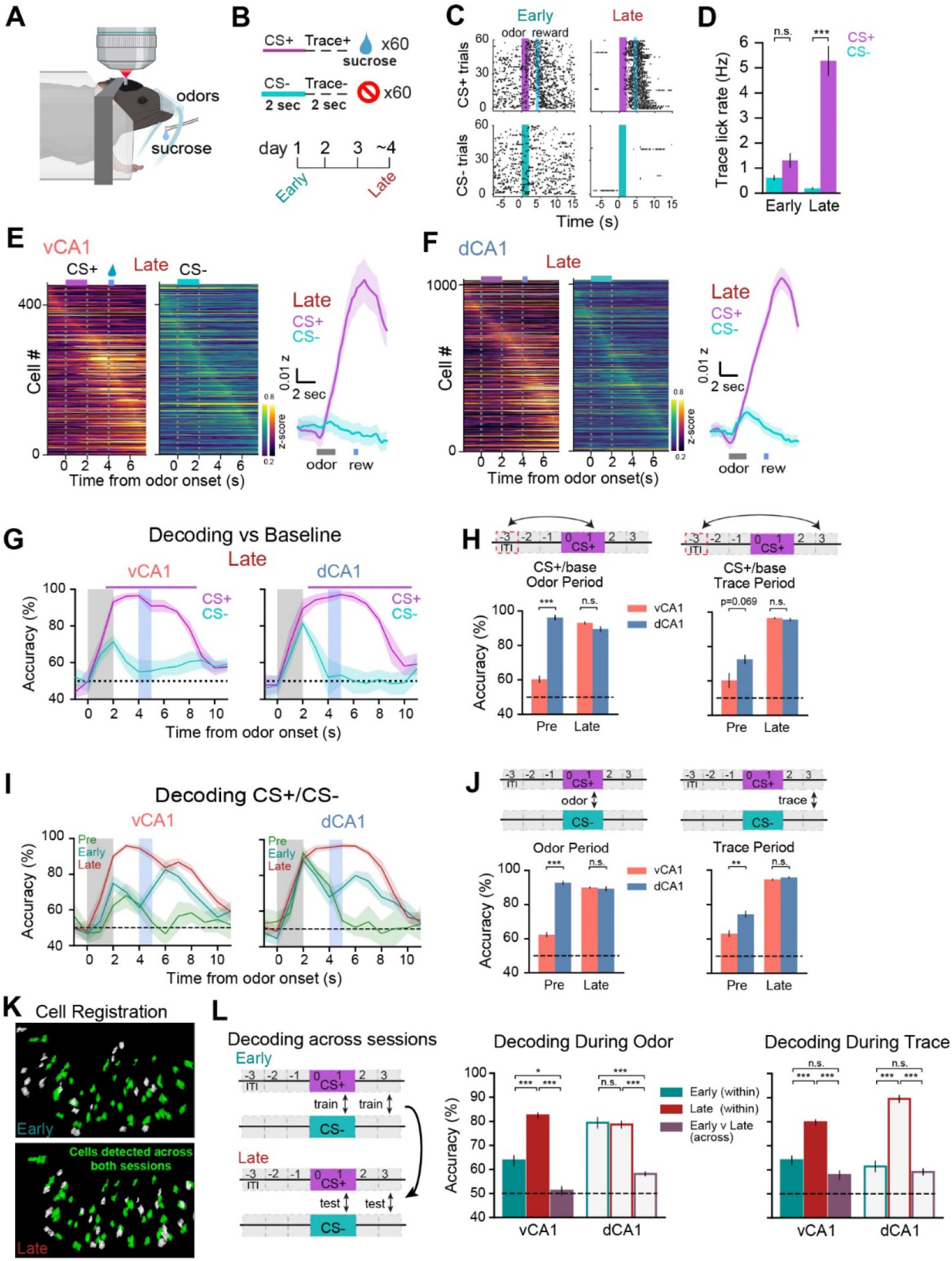
Discrimination training enhances odor representations in vCA1, trace period representations in dCA1 and vCA1. (A and B) Task schematics. (C) Lick rasters for an individual animal. Black tick = lick. During the first session of training (Early) licking is unstructured but becomes restricted to the time periods directly before and after reward delivery following CS+/CS-discrimination learning (Late). (D) Mean lick rates during the 2-second pre-reward (trace) period for all animals (±SEM, Mann-Whitney U test). (E) (left) Mean z-scored fluorescent signals for all vCA1 cells during Late session, ordered by peak time bin. See Figure S2A for Early session. (right) Population mean (±SEM). (F) Same as in (E), but for dCA1. (G) Decoding accuracies for each trial type vs ITI baseline (±SD). Color-coded bar above shows periods where the corresponding trial type accuracy is significantly greater than the opposing trial type (p < 0.01, Mann-Whitney U test). (H) Decoding accuracies for CS+ vs baseline during odor (left) or trace (right) periods (±SEM, Mann-Whitney U test). (I and J) Same as (G) & (H) but decoding for trial type. (K) Example of cells from the same FOV registered across Early and Late sessions. (L) Across-session trial type decoding accuracies (±SEM, Mann-Whitney U test). Note that only cells tracked across Early and Late sessions were used for within-session decoding.

In vCA1, learning of the odor-reward association was accompanied by an overall increase in mean activity during the CS+ odor presentation (Figures 2E, S2B, S2C) and heightened ability to decode CS+ odor period activity from baseline activity (Figures 2H, S2D), suggesting that assigning value to a stimulus leads to increased odor-evoked activity and encoding in ventral CA1. In addition, odor classification accuracy greatly increased in Late sessions after learning when compared with Pre and Early sessions (Figures 2I, 2J, S2E), indicating CS+/CS-representations become more distinct in vCA1 with learning. This was in contrast to dCA1, where odor decoding accuracy was high prior to training and remained so with learning (Figure 2G-2J, S2D, S2E).

As odor identities were reliably represented in dCA1 before training, we asked whether representations remained stable over the course of training. For this, we applied a cross-session classifier to neurons tracked across sessions (Figure 2K), training with data from the Early session and testing classification accuracy using data from the Late session (and vice-versa). As expected from our within-session results, in vCA1 odor representations changed with learning. However, in dCA1 surprisingly we found that odor representations also differed across training sessions, as cross-session decoding of CS+/CS-during the odor period was significantly worse relative to within-session decoding (Figure 2L). Therefore, although CS+/-representations are highly separable both before and after training, the representational geometry of CS+/-odors in dCA1 is altered with learning.

We next examined representations in the trace period, during which learned animals anticipate reward availability. We found parallel changes in trace-period representations in dCA1 and vCA1 with learning (Figures 2E-2J, S2B-S2E). Mean trace-period-evoked activity in both dCA1 and vCA1 markedly increased following CS+ delivery, but not CS-delivery. Correspondingly, CS+/CS-trial-type decoding accuracy during the trace period was significantly higher in both regions following learning. These trace-period representations emerged in concert with the initial signs of behavioral learning (Figures S2F, S2G), could not be explained solely by licking behavior (Figure S2H), and were distinct from odor-period representations (Figure S2I).

Together, these data suggest that a representation of odor identity is present in dCA1, independent of learning, while odor representations in vCA1 show greater dependence on learned behavioral significance. In addition, with learning both vCA1 and dCA1 are recruited during the trace period prior to reward delivery, seemingly encoding information related to the expectation of reward.

### Learned representations of task elements are modality-independent and learning-dependent

We next determined whether our results in CA1 generalized to other stimulus modalities, and whether neuronal changes that emerged with training were indeed learning dependent. For this, we trained a separate cohort of mice on a more difficult auditory cue discrimination task (Figure 3A, 3B), where an auditory cue (CS+) and a sucrose reward were separated by a 2s trace period, while a distinct CS-auditory stimulus was unrewarded. Reward delivery was contingent on licking during a 2-second reward availability window directly following the trace period. Unlike the odor-based task, which all animals quickly learned, mice took longer to learn this task (11.7 ± 1.9 days) with some failing to learn altogether (Figures 3C and S3E).

**Figure 3.**
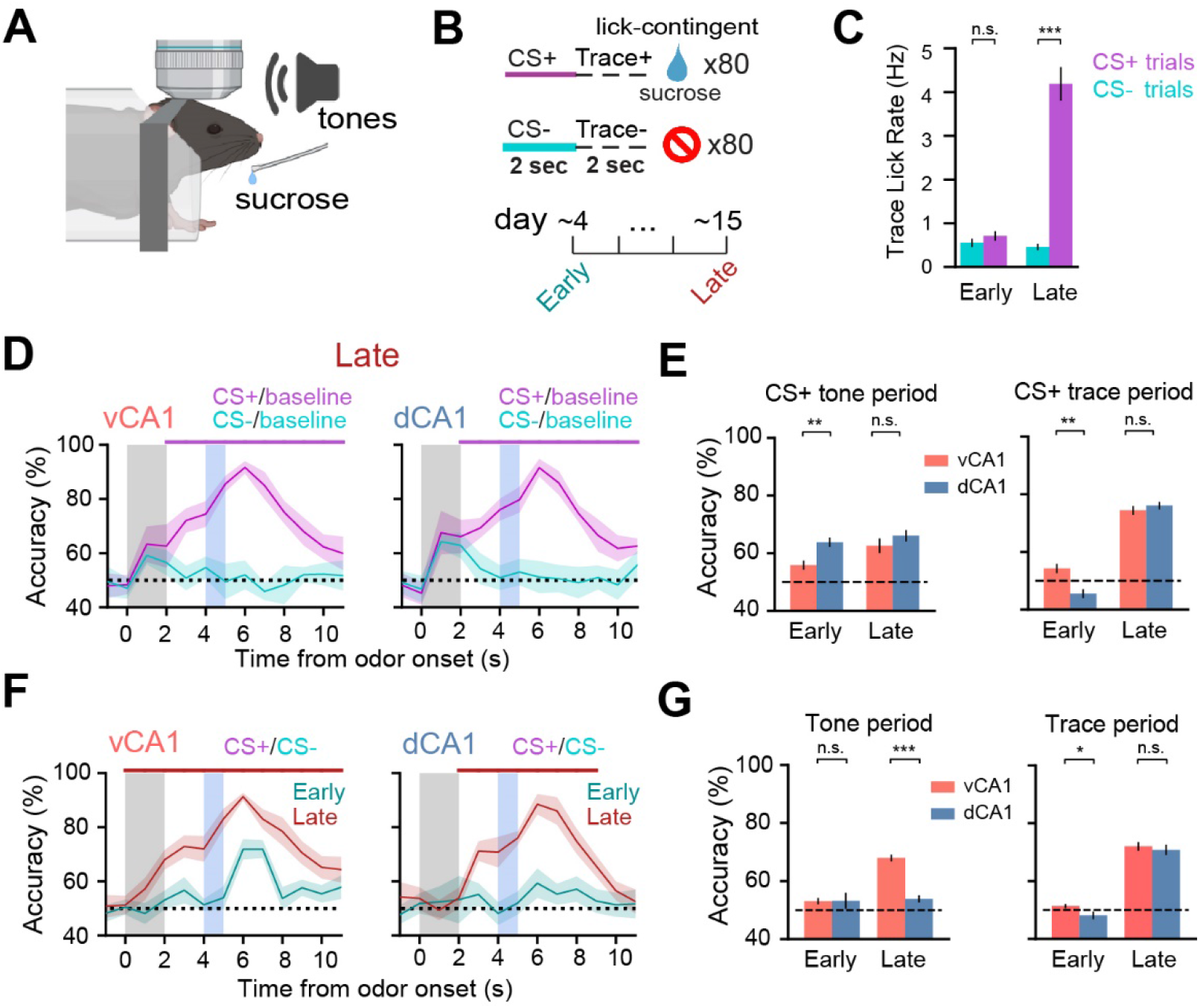
Learned representations of task elements are modality-independent and learning-dependent. (A and B) Task schematics (C) Mean lick rates during the 2-second pre-reward (trace) period for all animals (±SEM, Mann-Whitney U test). (D) Decoding accuracies vs baseline (±SD). Color-coded bar above shows periods where the corresponding trial type accuracy is significantly greater than the opposing trial type (p < 0.01, Mann-Whitney U test). (E) CS+ vs baseline decoding accuracies during odor (left) or trace (right) periods (±SEM, Mann-Whitney U test). (F and G) Same as (D) and (E), but decoding trial type.

As with the olfactory-based task, CS+ activity during tone presentation was more accurately classified from baseline activity in vCA1 after learning (Figures 3D, 3E, S3C), and classification of CS+/CS-tone identities likewise improved with learning (Figures 3F, 3G). This suggests that CS+ and CS-representations become more distinct in vCA1 over the course of discrimination training, regardless of the CS modality. In line with the odor-based task, we also found that decoding of CS+/CS-during the trace period was improved with learning in both regions (Figure 3D-3G).

Interestingly, although the CS+ and CS-tones could each be decoded from baseline activity with moderate accuracy in dCA1 (Figure 3D, 3E), tone identity could not be decoded accurately from dCA1 population activity either before or after learning (both ∼50% accuracy; Figure 3F, 3G). This contrasts with odorant identities, which could be consistently decoded with high accuracy, and is likely due to the greater ethological salience of odors vs tones. Despite this lack of strong encoding of tones in dCA1, these results are consistent with our odor task in that learning enhances the separability of cues in vCA1, but not dCA1.

A subset of vCA1 mice (n=3) failed to learn the tone-based version of the task, even after 20 days of training. In these “nonlearners” we found classification accuracy of CS+/CS-trial type during the tone or trace periods did not change between Early and Late sessions (Figure S3F, S3G). Thus, representations that emerge with learning are not simply driven by repeated exposure to task stimuli.

### Learned odor representations in vCA1 are sensitive to extinction but can be rapidly reinstated

Collectively, these results suggest that imbuing a stimulus with value enhances its representation in vCA1. How might stimulus representations change upon extinction of the odor-reward contingency? Would vCA1 continue to exhibit strong representations of odor, perhaps reflecting an enduring memory of the CS-US association, or would decoding performance fall back to baseline levels, suggesting vCA1 signals track current stimulus value?

After mice learned the odor-reward association, we extinguished it, omitting reward from CS+ trials. Mice rapidly extinguished conditioned responding early in the first session of extinction (Figure 4A-4C). In an extinction retrieval session 24 hours later, we found that odor classification accuracy resembled that of early, pre-learning sessions; that is, low in vCA1 but high in dCA1 (Figure 4D, 4E, S4A, S4B).

**Figure 4.**
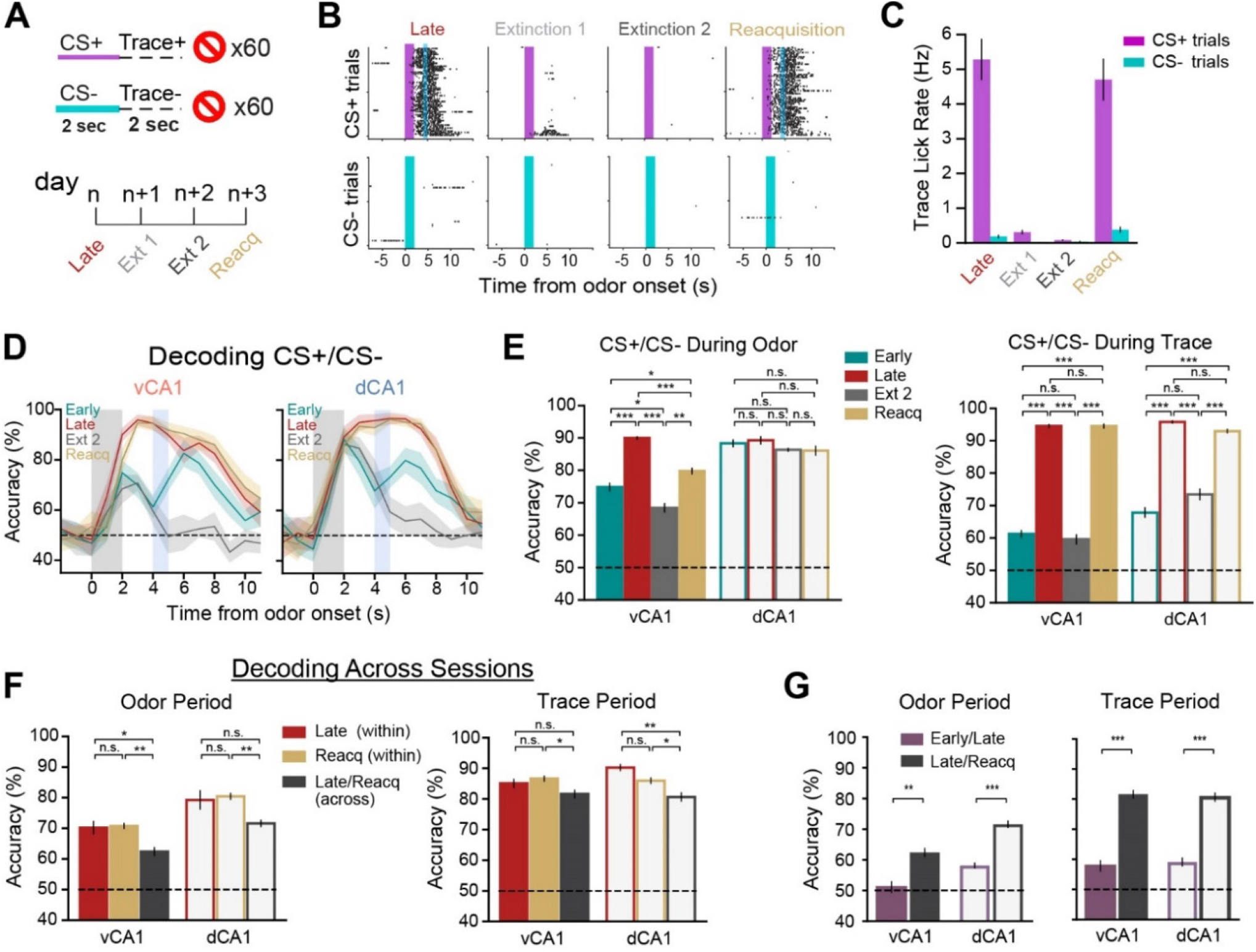
Learned odor representations are sensitive to extinction but can be rapidly reinstated. (A) Task schematic. (B) Lick rasters from an individual animal. Note the near absence of licking during the second day of extinction training and rapid resumption of anticipatory licking during the reacquisition session. (C) Mean trace-period lick rates (±SEM) across all animals. (D) Trial-type decoding accuracies (±SD). Early and Late are as in Figure 2I and are shown for reference. (E) Trial-type decoding accuracies during odor (left) or trace (right) periods (±SEM, Mann-Whitney U test). (F) Across-session trial-type decoding accuracies (±SEM, Mann-Whitney U test). Note that only cells tracked across sessions were used for within-session decoding here. (G) Decoding accuracy across sessions is significantly higher following learning, suggesting stable population codes during odor and trace periods.

The following day we reinstated conditioned responses in a reacquisition session. Animals rapidly resumed anticipatory licking behavior during CS+ trials, indicating an intact memory of the rewarded task structure. Correspondingly, odor identity classification accuracy increased in vCA1 (Figures 4D, 4E, S4A, S4B). These data indicate that the discriminability of odor representations in vCA1, but not dCA1, is sensitive to the current value associated with an odor.

To investigate whether representations were stable before and after the extinction sessions, we used a cross-session classifier using data from cells tracked across Late and Reacquisition sessions. We found that odor-and trace-period representations remained relatively stable across extinction in both vCA1 and dCA1, as trial type could be accurately classified across sessions during both periods (Figure 4F). This contrasts with the instability of odor representations observed during initial learning (Figures 2, 4G, S4D, S4E) and indicate that, once learned, representations of odor and reward anticipation are to a large extent stable across days and across the degradation and reinstatement of odor-reward contingencies, indicating that CA1 may be a storage site for these representations

### Long-timescale representations in vCA1

In vCA1, place fields are broader than those in dHPC, which has been hypothesized to allow vCA1 to contain global representations of behaviorally relevant contexts (Chockanathan and Padmanabhan, 2021; Harland et al., 2017; Jung et al., 1994; Kjelstrup et al., 2008). We likewise examined whether vCA1 contained more diffuse representations than dCA1 in our task as the mouse “moved” through the trial. We trained a linear classifier to discriminate trial type (CS+ vs CS-) using data from a single time bin, then tested classification accuracy on every other time bin (Figure 5A). We found that in vCA1, but not dCA1, there was a persistent trial-type representation throughout the trial duration, and trial-type could be decoded even when training and testing on time bins separated by +/-5s seconds (Figures 5B, 5C). Importantly, this was only observed for time bins within the trial duration (1s post odor onset through 4s post reward delivery), and most prominently for sessions where the CS-US contingency had been learned and was actively being rewarded. As these prolonged representations in vCA1 may in part arise from protracted individual neuron activity in vCA1 compared to dCA1, we examined the distribution of activity in these regions during trials. Indeed, we found that vCA1 displayed broader firing within trials than dCA1 (Figures S5A-B).

**Figure 5.**
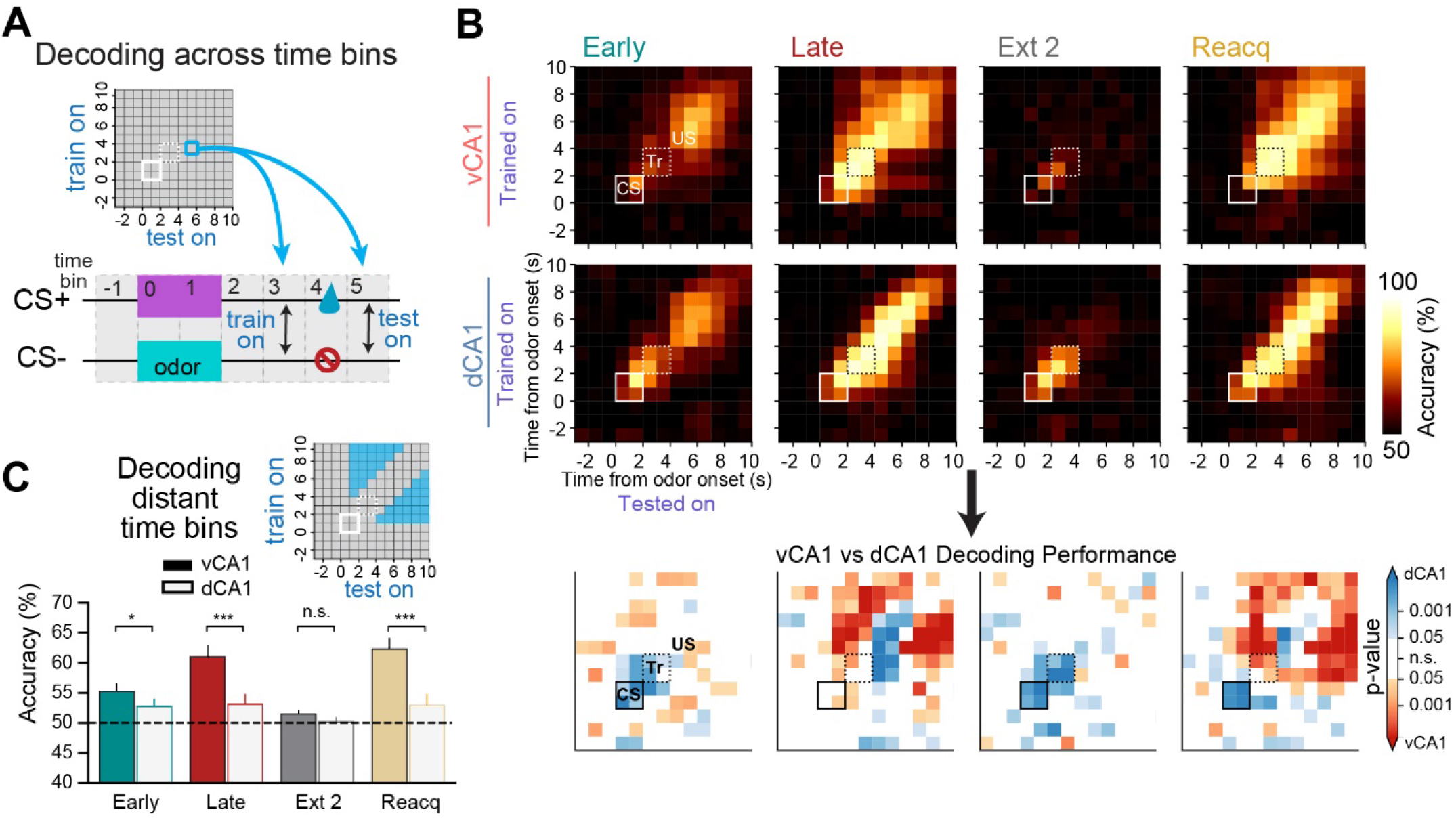
Long timescale representations in vCA1. (A) A linear classifier was trained to discriminate trial type using data from a single time bin, then tested for decoding accuracy on all other time bins. Each square is the decoding accuracy for the corresponding time bins on the x- and y-axes. The blue square shows the result corresponding to training the classifier on activity three seconds post odor onset and testing five seconds post odor onset. (B) Decoding across time bins during different sessions of the 2-odor task. Closed white square denotes odor period (CS). Dashed square denotes trace period (Tr). Matrix at bottom reports p-values comparing the decoding accuracies of vCA1 and dCA1 (Mann-Whitney U test). (C) Analysis of cross-time-bin decoding results for comparisons separated by at least 2 time bins (±SEM, Mann-Whitney U test). Inset: blue filling shows data bins that were included for analysis.

### Pre-reward signals generalize across distinct predictive cues

Our results thus far show that, with learning, both hippocampal regions display strong representations during the odor and trace periods that are largely distinct from one another. To better understand what information is being encoded during these periods, we trained mice with four odor stimuli; two that were always followed by sucrose reward (CS1+ and CS2+), and two that were followed by no outcome (CS3- and CS4-). This design allowed us to directly test the similarity of representations across trial types with distinct cue identities but the same outcome (Figures 6A, 6B).

**Figure 6.**
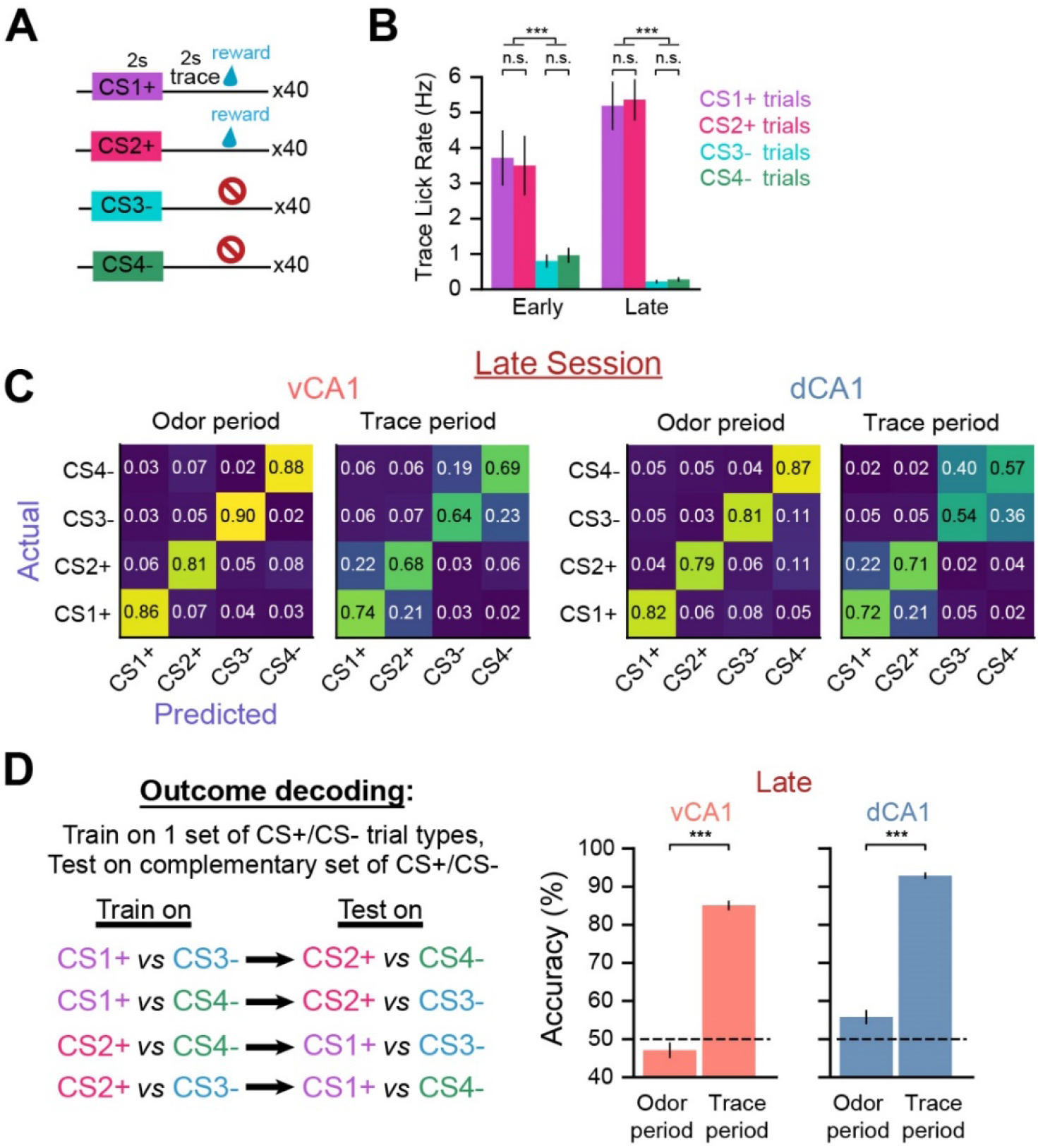
Pre-reward signals generalize across distinct predictive cues. (A) 4-odor task schematic (B) Mean trace-period lick rates across all animals (±SEM, Mann-Whitney U test). (C) Confusion matrices for decoding trial type from population activity. The y-axis denotes the actual trial type experienced and the x-axis indicates the proportion of trials for which each trial type was decoded. The ascending diagonal represents the proportion of trials correctly classified, while other row entries indicate the proportion of trials where the actual state was confused with the corresponding trial type. Note the high accuracy during the odor delivery period and the increased incidence of parallel trial types (e.g., CS1+ and CS2+) being confused during trace. (D) A linear classifier was trained to discriminate activity between reward-predictive and non-predictive trial types (e.g., CS1+ vs CS3-), then tested using data from the complementary trial types (CS2+ and CS4-). The mean for all combinations of trial-type pairs is presented (±SEM, Mann-Whitney U test).

We first tested how well a linear classifier could predict each of the four trial types using population activity during the odor or trace periods (Figure 6C). We found that, following learning, odor identity could be predicted with high accuracy during the odor-delivery period for both dCA1 and vCA1. Conversely, although individual trial types remained discriminable during the trace period, classification accuracy was lower in both regions during this period, with classifier errors predominantly occurring between trial types with the same outcome (e.g., CS1+ and CS2+). Thus, after learning, trace period activity is highly discriminable between trial type categories (CS+ vs CS-), but less so within each category.

The reduced decoding accuracy between CS1+ and CS2+ trial types during the trace period suggests a common signal across these trials. To more directly test this, we trained a linear classifier to discriminate activity between a reward-predictive trial type (e.g., CS1+) versus a non-predictive trial type (e.g., CS3-), then tested classification accuracy using data from the complementary trial types (CS2+ and CS4-), which the decoder had never seen (“outcome decoding”; see Figure 6D). Here, high decoding accuracy would indicate similar neural states across related trial types (e.g., CS1+ and CS2+).

After learning, we found high outcome decoding accuracy during the trace period in both dCA1 and vCA1 (Figure 6D), further indicating that there exists a representation related to reward expectancy that is independent of the identity of the stimulus that precedes it. In contrast, outcome could not be decoded during the odor delivery period, suggesting that CS identity or specific CS-outcome associations are primarily represented during odor delivery. Dimensionality reduction analysis returned results that were analogous with the above (Figure S6). Together, these data indicate a switch from individual odor representations (during odor delivery) to representations related to expected outcome (during trace) as animals progress through a trial.

### Aversive conditioning and reversal learning

We next determined whether our results from appetitive learning generalize to aversive conditioning. For this, we trained mice in an associative learning task with three novel CS odors (Figure 7A) that were paired with either sucrose (CS+rew), tail shock (CS+sh), or nothing (CS-). In general, our CS+sh results were qualitatively similar to those for the CS+rew condition, and both conditions mimicked simple (2-odor) appetitive learning (Figures 7 and S7). Of particular note, however, trace period decoding accuracy was significantly lower prior to shock compared to reward (Figures 7C - 7F, S7A - S7C).

**Figure 7.**
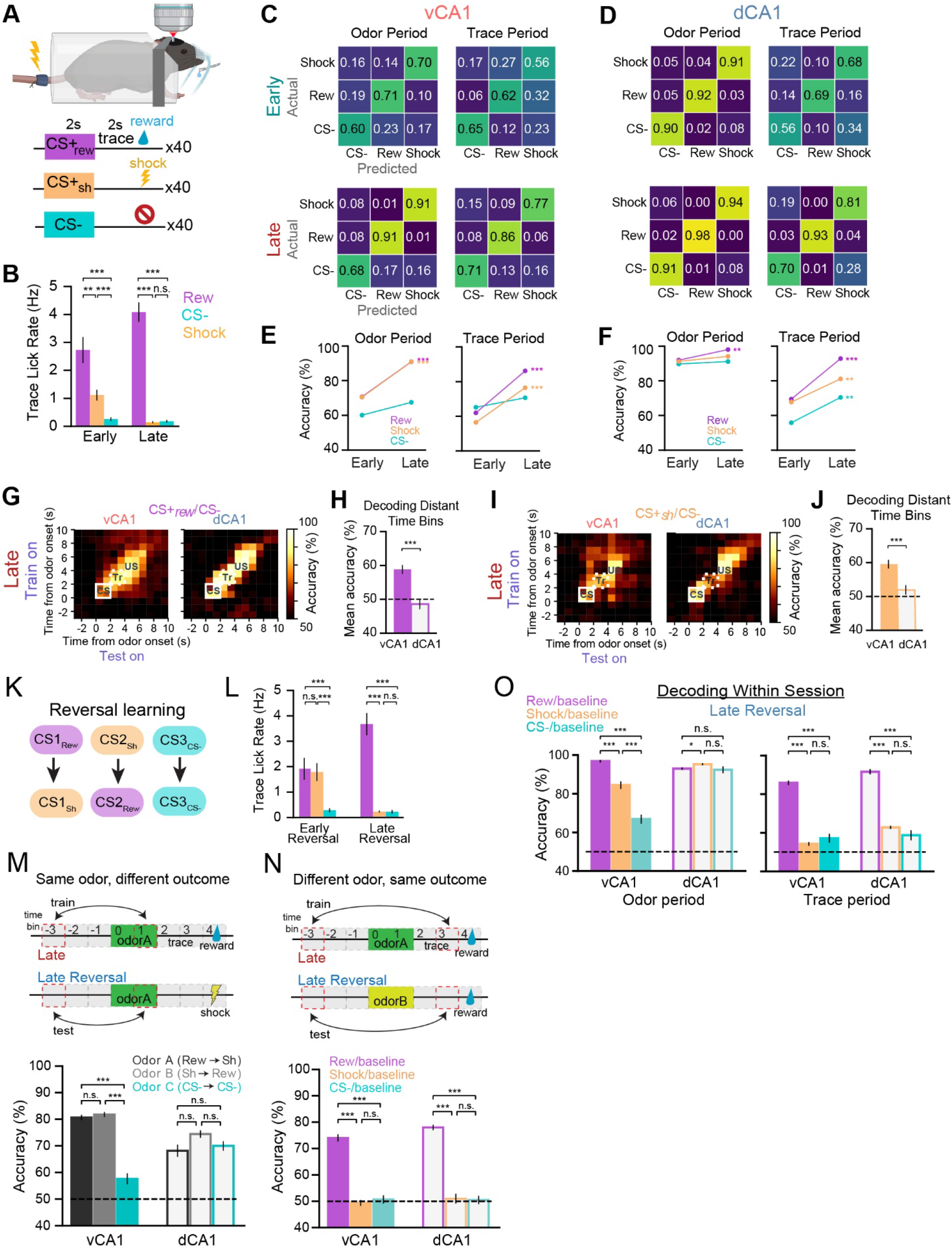
Aversive conditioning and reversal learning. (A) Task schematic. (B) Mean trace-period lick rates (±SEM, Mann-Whitney U test). (C and D) Confusion matrices for trial-type decoding accuracy during Early (upper) or Late (lower) sessions. (E and F) Line plots of results in (C) and (D), respectively. Statistics compare Early vs Late decoding accuracies (±SEM, Mann-Whitney U). (G) Decoding CS+reward vs CS-trials across time bins during Late. (H) Analysis of cross-time-bin decoding results for comparisons separated by at least 2 time bins (±SEM, Mann-Whitney U test). Inset: blue filling shows data bins that were included for analysis. (I and J) Same as (G), (H) but decoding for CS+shock vs CS-trials. (K) Mice were trained on reversed CS-US contingencies, while CS-trials remained unchanged. (L) Mean trace-period lick rates across all animals (±SEM, Mann-Whitney U test). (M) (upper) Schematic showing cross-session odor vs baseline decoding for a specific odor paired with different outcomes. (lower) Cross-session decoding accuracies (±SEM, Mann-Whitney U test). (N) Same as in (M) but decoding during the trace period for a specific outcome preceded by different odors. (O) Within-session decoding accuracies for each trial type vs baseline during odor (left) or trace (right) periods (±SEM, Mann-Whitney U test).

Thus far, our data have not resolved whether odor-period representations in trained animals reflect odor identities or perhaps information more representative of memory, such as specific cue-outcome associations. To adjudicate between these possibilities, after training mice to associate one odor with sucrose (CS+rew) and another with shock (CS+sh), we reversed the contingencies, where the previously rewarded odor was now paired with shock and vice-versa (Figure 7K). Animals were trained on the new contingency until anticipatory licks were only observed during the CS+rew trial (Figure 7L), and neurons were tracked across all sessions. To probe for stable encoding of odor identity across different US pairings, we decoded across reversal learning, training a linear classifier with population data from the final session prior to reversal (Late), and tested classification accuracy using data after the reversed contingencies had been learned (Late Reversal). In both dorsal and ventral CA1, we found that the neural representation for a specific odor remained intact regardless of whether the odor predicted sucrose or shock (Figures 7M, S7G).

Finally, we assessed whether outcome expectation signals during the trace period remained stable following reversal learning, and whether stability differed for reward vs. shock outcomes. For this, we performed the same analysis as above, but using trace period data. In both dCA1 and vCA1, the cross-session decoder performed well when discriminating reward anticipation signals preceded by different odors (Figures 7N, S7H), suggesting a conserved signal related to reward anticipation that is independent of the odor that precedes it. Conversely, there was not a conserved representation across reversal learning when anticipating shock. Additional analyses revealed this was due to the absence of pre-shock trace-period signals during the Late Reversal session (Figure 7O). These results indicate that odor representations in both dCA1 and vCA1 are independent of the nature of the associated US, and that stable signals preceding reward, but not shock, emerge with learning in these regions.

## DISCUSSION

### Relevance-dependent stimulus encoding in vCA1

Prior to cue-reward training, we found that dCA1 strongly encodes the identity of individual odors, in line with previous findings showing that environmental stimuli need not be paired with reward or other salient outcomes to be represented in dCA1 (Li et al., 2017; Taxidis et al., 2020). In contrast, vCA1 was only weakly reactive to odors prior to training, and odor identity was poorly decoded using population activity. Instead, odor decoding was generally contingent on salience, increasing for odors predictive of salient outcomes (e.g., reward or shock), and decreasing in the absence of these outcomes (e.g., extinction). These data extend previous findings showing the dorsal and intermediate HPC intensify activity for stimuli and locations with learned significance (Eichenbaum et al., 1987; Jin and Lee, 2021), and further show that these changes manifest as distinct patterns of population activity that effectively decouple stimulus representations.

Although odor representations in vCA1 only emerged when an odor gained predictive value, it is important to note that vCA1 does not represent value per se, as population activity differed for distinct odors with the same associated value. We also did not find evidence that the associated outcome was represented by population activity during odor delivery, although it’s possible this information was represented by a small subset of cells whose signal was too weak to appreciably influence the decoder. Instead, our data suggest vCA1 strongly encodes the identity of stimuli that are relevant to the animal. This dependency on relevance is fundamentally different from that observed in dCA1, where the separability of stimulus representations was not dependent on their perceived relevance. Such relevance-selective processing in vCA1 may be important for conveying stimulus-value information to the frontal cortex (Wikenheiser and Schoenbaum, 2016; Burton et al., 2009), passing information to emotional centers, such as the amygdala, for further processing (Felix-Ortiz et al., 2013; Graham et al., 2021; Xu et al., 2016), or alerting downstream mediating approach/avoidance behaviors (LeGates et al., 2018; Trouche et al., 2019).

### Stability and dynamism of odor representations in CA1

Following learning, odor representations remained relatively stable across sessions, including across extinction training or when the valence of the paired outcome was switched, indicating CA1 is a storage site for odor representations with learned relevance. Of note, however, across-session decoding was consistently lower than within-session decoding, suggesting that HPC representations are continually updated. Dynamic representations could reflect differences in the external (time, location) or internal (motivation, attention) context in which an event is experienced, properties which are known to affect HPC processing (Cai et al., 2016; Kennedy and Shapiro, 2009; Mankin et al., 2012; Radvansky et al., 2021; Ziv et al., 2013) and may provide a neural mechanism for differentiating distinct (novel contexts), yet related (same odor), experiences.

### Encoding during outcome anticipation across the dorsoventral axis

In dCA1, neuronal activity has been shown to be modulated in rewarded tasks through various means, including accumulation of place fields at rewarded locations (Danielson et al., 2016; Dupret et al., 2010; Kaufman et al., 2020; Sato et al., 2018; Xu et al., 2019) and, in tasks where goal location is dissociated from reward, an increase in out-of-field firing at the goal (Duvelle et al., 2019; Hok et al., 2007). Further, dedicated populations of goal-approach cells have been identified across the dorsoventral axis (Ciocchi et al., 2015; Eichenbaum et al., 1987; Gauthier and Tank, 2018; Markus et al., 1995; Royer et al., 2010). Our results extend these findings beyond the spatial domain and show that ventral CA1 also contains a dedicated signal during reward anticipation that generalizes across predictive cues and is stable across days.

Most findings observed with appetitive conditioning were likewise seen with cue-shock conditioning. Surprisingly, however, shock anticipation was only weakly encoded by both dCA1 and vCA1, despite robust activation of both regions in response to cue and shock deliveries. Moreover, shock anticipation signals were further diminished with subsequent training, eventually becoming indistinguishable from ITI activity. Although previous reports examining anticipation of aversive stimuli are mixed for dCA1 (Ahmed et al., 2020; MacDonald et al., 2013; Mount et al., 2021; Zhang et al., 2019), this finding is particularly surprising for vCA1, which is known to mediate anxiety and fear responses (Jimenez et al., 2020; Kjelstrup et al., 2002).

Why would the hippocampus strongly represent the expectation of reward, but not shock, particularly in light of the crucial role the HPC plays in trace fear conditioning (Bangasser et al., 2006; McEchron et al., 1998)? One important difference between our reward and shock paradigms is that reward is contingent on behavior (licking) whereas shock is inescapable and thus behaviorally irrelevant. Interestingly, VTA dopamine (DA) neurons reduce their responding to aversive stimuli following repeated exposure to inescapable shocks (Belujon and Grace, 2014; Wu et al., 2021). As dopamine signaling is believed to play a key role in hippocampal plasticity (Palacios-Filardo and Mellor, 2019), it is possible this reduction in DA signaling may contribute to the loss of CA1 shock expectation encoding and would also suggest that DA signaling is continually required to maintain outcome-expectation signals.

Considering the dorsoventral differences in CS encoding with learning, it is interesting to note the parallel changes in these regions when anticipating specific outcomes, although perhaps not surprising. For example, both regions play a vital role in reward learning, particularly via their projections to the nucleus accumbens (NAc) (Britt et al., 2012; LeGates et al., 2018; Trouche et al., 2019). Recent evidence, however, shows dCA1 and vCA1 differentially engage the NAc (Sosa et al., 2020), suggesting these regions convey distinct information to downstream targets. The nature of that information is unclear, as signals observed during the trace period may be conveying the expectation of a specific outcome, promoting specific behaviors (e.g., approach or behavioral inhibition) and/or conveying a more general signal of motivational salience or valence. While our study was not designed to adjudicate these possibilities, the presence of distinct representations during anticipation of reward or shock (Figs 7C-7D) argues against a general arousal signal during this time. Additionally, the lack of pre-shock signals following extended training suggests negative valence is likely not encoded in either region during this time.

### Dorsal and ventral CA1 multiplex task-related information

Consistent with previous reports (Bernardi et al., 2020; McKenzie et al., 2014; Nieh et al., 2021; Stefanini et al., 2020) we find that dCA1 multiplexes information. Specifically, we find that during the trace period dCA1 conveys information relevent to expected outcome simultaneous with the identity of the preceding odor. We now show vCA1 similarly multiplexes this information. Consequently, a downstream recipient of these signals could not only decode whether reward is forthcoming, but at the same time the identity of the cue that preceded it, which may be important for updating cue value. Alternatively, it is possible that outcome and cue identity signals present during the trace period are each routed to distinct downstream targets (Beyeler et al., 2016; Namboodiri et al., 2019; Otis et al., 2017), consistent with vCA1 circuitry, where specialized functions are parsed across vCA1 projection pathways (Ciocchi et al., 2015; Jimenez et al., 2018, 2020; Shpokayte et al., 2020; Xia and Kheirbek, 2020; Xu et al., 2016).

### Extended population representations in vCA1

Similar to differences in single-cell place field properties across the DV axis, where dCA1 spatial representations are more circumscribed and vCA1 representations generalize across large swaths of space (Jung et al., 1994; Kjelstrup et al., 2008), our cross-time-bin population analysis suggests a more rapid turnover of population activity patterns (neural states) in dCA1 compared to vCA1. Thus, the broad firing observed in vCA1 during spatial exploration may reflect a more general property of this region that extends beyond representations of physical space. These results are also in line with human studies of memory that suggest posterior HPC is associated with recall of detailed information, such as the temporal sequence of events, while anterior HPC represents higher level information, such as the location of where the collection of events occurred (Harland et al., 2017; Poppenk et al., 2013).

This longer timescale of neural representations observed in vCA1 may serve to link discontinuous cue-reward events, providing a neural substrate through which credit can be assigned to the stimuli or actions that preceded reward delivery (Petter et al., 2018; Sosa and Giocomo, 2021; Stachenfeld et al., 2017). Given that representations in vCA1 stretched across the duration of a trial, these signals may alternatively serve to “locate” the animal within the task space (Knudsen and Wallis, 2021), such as the task context (i.e., trial type) currently being occupied, and broadcast this information to downstream regions (e.g., frontal cortex) to retrieve context-relevant memories and guide behavior (Komorowski et al., 2013; Wikenheiser and Schoenbaum, 2016; Wikenheiser et al., 2017).

### Mechanisms underlying changes in neural representations in CA1

Our results show vast reorganization of hippocampal activity networks during associative learning. Unknown, however, are the mechanisms responsible for implementing this change. Dopaminergic, cholinergic, serotonergic, and adrenergic signals are all present to varying degrees across the DV axis of the hippocampus (Basu and Siegelbaum, 2015; Palacios-Filardo and Mellor, 2019), and each is integral for hippocampal plasticity and learning (Gu and Yakel, 2011; Kaufman et al., 2020; McNamara et al., 2014; Palacios-Filardo and Mellor, 2019, 2019; Teixeira et al., 2018). Dopamine signaling, for one, mediates reward-dependent reorganization of place fields (McNamara et al., 2014) and may analogously promote changes in CS+ representations by integrating stimulus identity signals with reward-induced DA release.

In addition to modulatory inputs, the HPC receives information from a multitude of extrahippocampal areas that may further shape CA1 network activity. The orbitofrontal cortex (OFC) is postulated to provide the hippocampus with information about expected outcomes (Wikenheiser and Schoenbaum, 2016), and may contribute to the reward-anticipation signals seen here. Input from the medial thalamus and/or amygdala to vCA1 (Gergues et al., 2020) could provide additional information regarding the learned salience or valence of stimuli (Beyeler et al., 2016; Felix-Ortiz et al., 2013; Ramanathan et al., 2018). Additionally, we recently showed that odor representations in lateral entorhinal cortex (LEC) become more separable with learning (Woods et al., 2020) and may thus influence changing odor representations in CA1 across training.

Finally, intrahippocampal signaling may also contribute to changes seen here. Although recurrent connectivity in CA1 is sparse (Deuchars and Thomson, 1996; Knowles and Schwartzkroin, 1981) (but see (Yang et al., 2014)), learning is known to augment recurrent interactions in the brain (Albieri et al., 2015; Biane et al., 2019) which can amplify inputs and induce attractor networks (Douglas and Martin, 2007; Lien and Scanziani, 2013), developments that could mediate the separation of odor representations we observed in vCA1. Additionally, inhibitory and astrocytic signaling may contribute to these changes (Bazargani and Attwell, 2016; Turi et al., 2019).

## Conclusions

Here we have shown that dCA1 and vCA1 are largely attuned to different aspects of the world. In simplest terms, the hippocampus might thus be thought of as undergoing a shift from externally based to internally based encoding of environmental variables along the DV axis. Such a division of labor could facilitate learning (Harland et al., 2017; Petter et al., 2018; Staresina and Davachi, 2009) and support the creation of a rich internal model that not only charts relationships in the world (dHPC), but also imbues certain relationships with meaning and emphasizes relevant stimuli (vHPC) (Collin et al., 2015; Harland et al., 2017; Shohamy and Wagner, 2008). Moreover, this simplified model is consistent with hippocampal lesion studies, where dHPC damage disproportionally affects declarative memory, and vHPC dysfunction is more closely associated with a failure to properly assign/update value, such as in PTSD, addiction, and depression (Fanselow and Dong, 2010; Strange et al., 2014).

On the other hand, we also saw a clear overlap in how some task variables are represented in dCA1 and vCA1. Why might these functionally distinct regions encode information that is seemingly redundant? As efferent connectivity patterns of dCA1 and vCA1 differ considerably (Bienkowski et al., 2018; Cenquizca and Swanson, 2007; Gergues et al., 2020; Strange et al., 2014), it is likely that each region is broadcasting this information to distinct downstream targets. When dCA1 and vCA1 outputs do converge onto the same region, these inputs may be handled differently, as appears to be the case with the NAc in reward learning (Sosa et al., 2020). Therefore, redundancy of representations across dorsal and ventral CA1 may be processed distinctly and uniquely influence ongoing operations. An interesting question for future inquiry is whether dCA1 and vCA1 inherit these overlapping representations from common or separate input source(s), or perhaps inform one another.

## Acknowledgements

We thank Vijay Namboodiri and Loren Frank for discussion and comments and Kenji Litke, Stephen Chien, Aadith Vittala, Aditya Garg and Clay Lacefield for technical assistance. JSB was supported by the Brain and Behavioral Research Foundation (NARSAD) and the Sandler PBBR Independent Postdoctoral Fellow Research Award. MAL was supported by NSF GR Fellowship. MAK was supported by NIMH (<u>R01 MH108623</u>, <u>R01 MH111754</u>, R01 MH117961), NIDCD (R01 DC019813) a One Mind Rising Star Award, a Research Grant from HFSP (Ref. No-RGY0072/2019), the Esther A. and Joseph Klingenstein Fund, the Pew Charitable Trusts, the McKnight Memory and Cognitive Disorders Award and The Ray and Dagmar Dolby Family Fund.

## Author contributions

Conceptualization, JSB, MAK, MAL; Methodology, JSB, MAK, MAL, NIW; Writing – Original Draft, JSB; Writing – Reviewing & Editing, JSB, MAK, MAL; Investigation, JSB, MAL, SPB, AF, SH, ND, DLA; Visualization, JSB, FS, MAK, MAL; Formal Analysis, JSB, FS, MAL, MAK; Supervision, MAK, JSB; Funding Acquisition, MAK, JSB.

## SUPPLEMENTAL FIGURES

**Figure S1.**
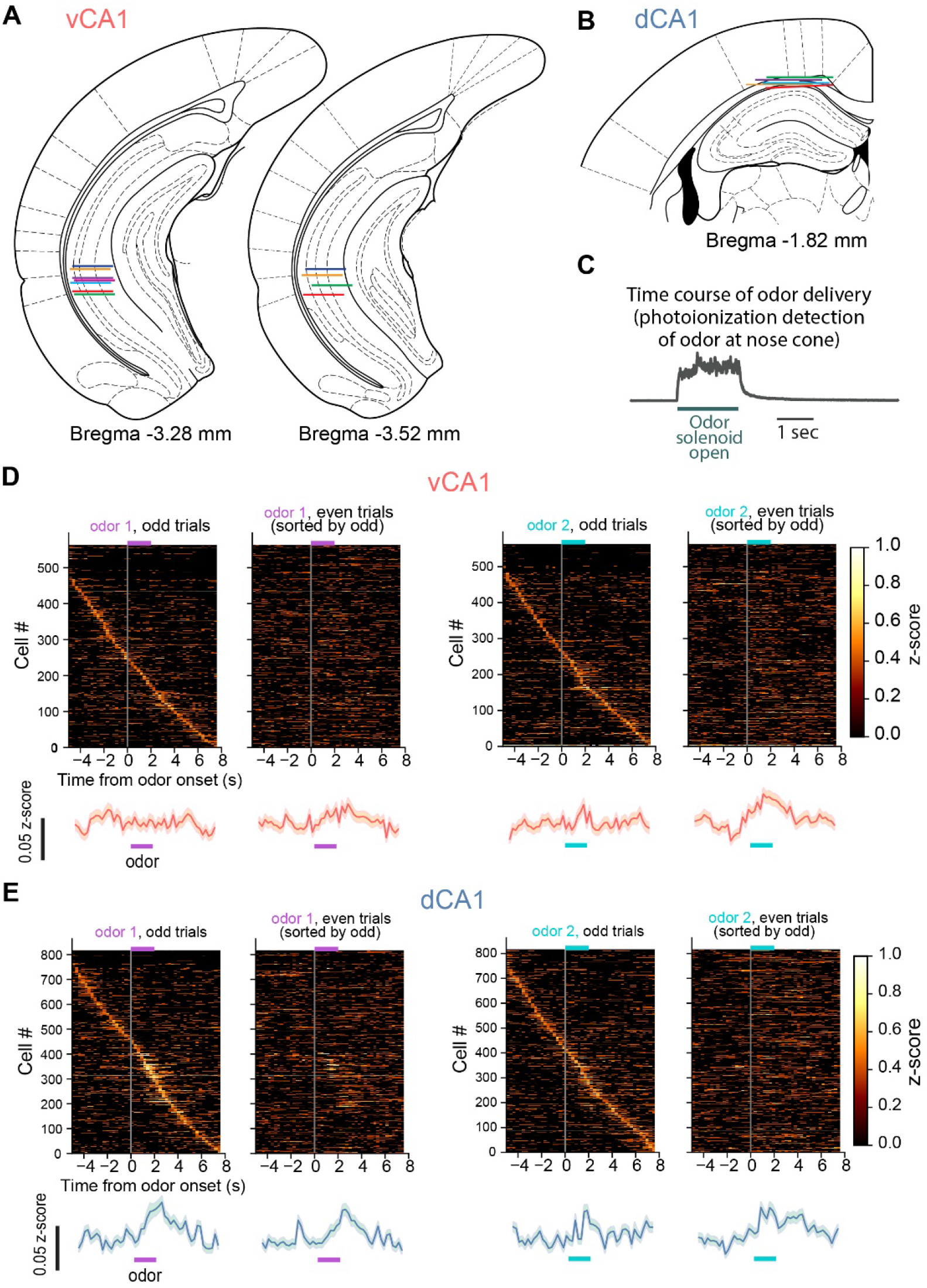
Related to Figure 1. Implant localization and pre-training neural activity. (A and B) Reconstructed GRIN lens implant locations for all vCA1 (A) and dCA1 (B) animals used in odor-based studies. Colored lines indicate the estimated location of the lens impression left on the tissue. Atlas images adapted from (Paxinos and Franklin, 2019). (C) Time course of odor presence at the nosecone. (D) Cross-validated neural activity during the Pre session. Each trial type (odor1 or odor2) was separated into odd and even trials, and vCA1 neural activity was z-scored. For each time bin, z-scores were averaged across all trial subsets, and sorted by peak firing rate latency during odd trials. Population mean is shown below (±SEM). (E) same as (D), but for dCA1.

**Figure S2.**
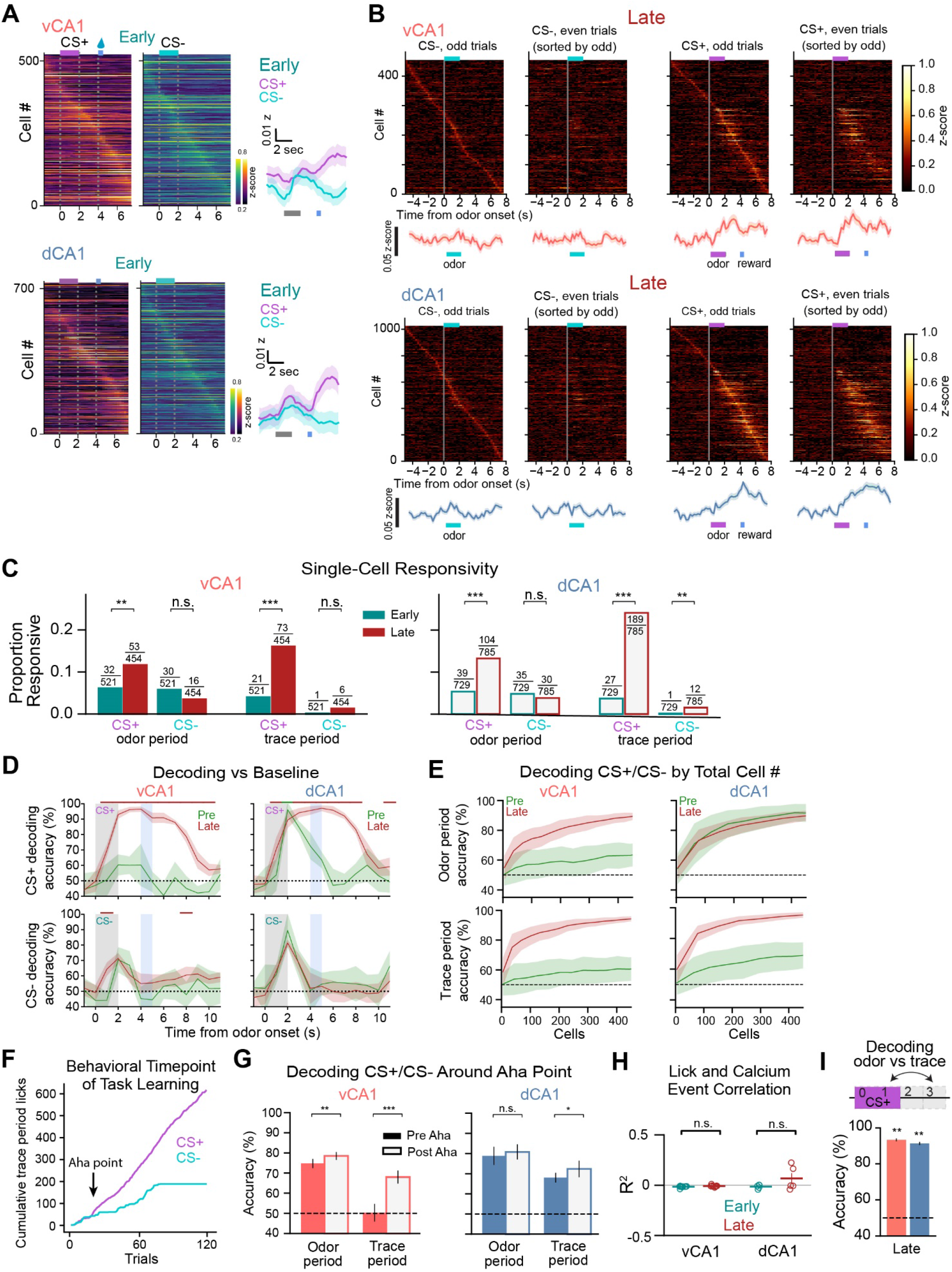
Related to Figure 2. Learning-related changes in neural activity and population decoding. (A) (left) Mean z-scored fluorescent signals for all recorded cells during the Late session, ordered by peak time bin. See Figures 2E and 2F for Late session. (right) Population mean (±SEM). (B) Cross-validated neural activity. Each trial type (CS+ or CS-) was separated into odd and even trials, and neural activity was z-scored. For each time bin, z-scores were averaged across all trial subsets, and sorted by peak firing rate latency during odd trials. Population mean is shown directly below heatmap (±SEM). (C) Proportion of neurons whose activity was significantly modulated during odor-or trace-period compared to pre-odor baseline (deemed “responsive cells”). Numerator denotes the number of responsive cells. Denominator denotes the total number of cells recorded. Fisher’s exact test. Statistical power for the pre-training session (Pre) was too low for meaningful analysis (only 15 trials/trial-type in Pre vs 60 trials/trail-type in Early and Late). (D) Population-activity decoding accuracy for CS+ or CS-trials from baseline (±SD). Color-coded bar above shows periods where the corresponding trial type accuracy is significantly greater than the opposing trial type (p < 0.01, Mann-Whitney U test). (E) Relationship between trial-type decoding accuracy and the total number of cells included in analysis (±SD). (F) Sample cumulative licking during the trace period for CS+ and CS-trials from the Early and second day of learning. The Aha point, in this example at trial 20, represents the first moment the difference between the cumulative licking in CS+ and CS-trials exceeded the learning threshold (see Methods). (G) Trial-type decoding accuracy during odor or trace periods using 30 CS+ and CS-trials before and after the Aha point. In vCA1, decoding accuracy significantly increases after the aha point for the odor and trace periods (p < .01 and p<.001, respectively, Mann-Whitney U test). Before the aha point, decoding during trace is not significantly different from chance (p = .88, Wilcoxon test). In dCA1, aha decoding does not significantly increase during the odor period and increases by a small but significant amount during trace period (p<.05, Mann-Whitney U test). dCA1 trace period decoding before the aha point is already significantly above chance (p < .01, Wilcoxon test). (H) Linear regression of lick rates and Ca2+ in vCA1 and dCA1 during Early and Late associative learning sessions (see Methods). We found that neural activity is not significantly correlated to lick rates (R^2^ is approximately zero for all animals in both sessions; t-test, p>0.05). (I) Decoding odor period vs trace period (±SEM). The high accuracy of decoding performance indicates there are different population activity states during the odor and trace periods (Wilcoxon signed-rank test vs chance).

**Figure S3.**
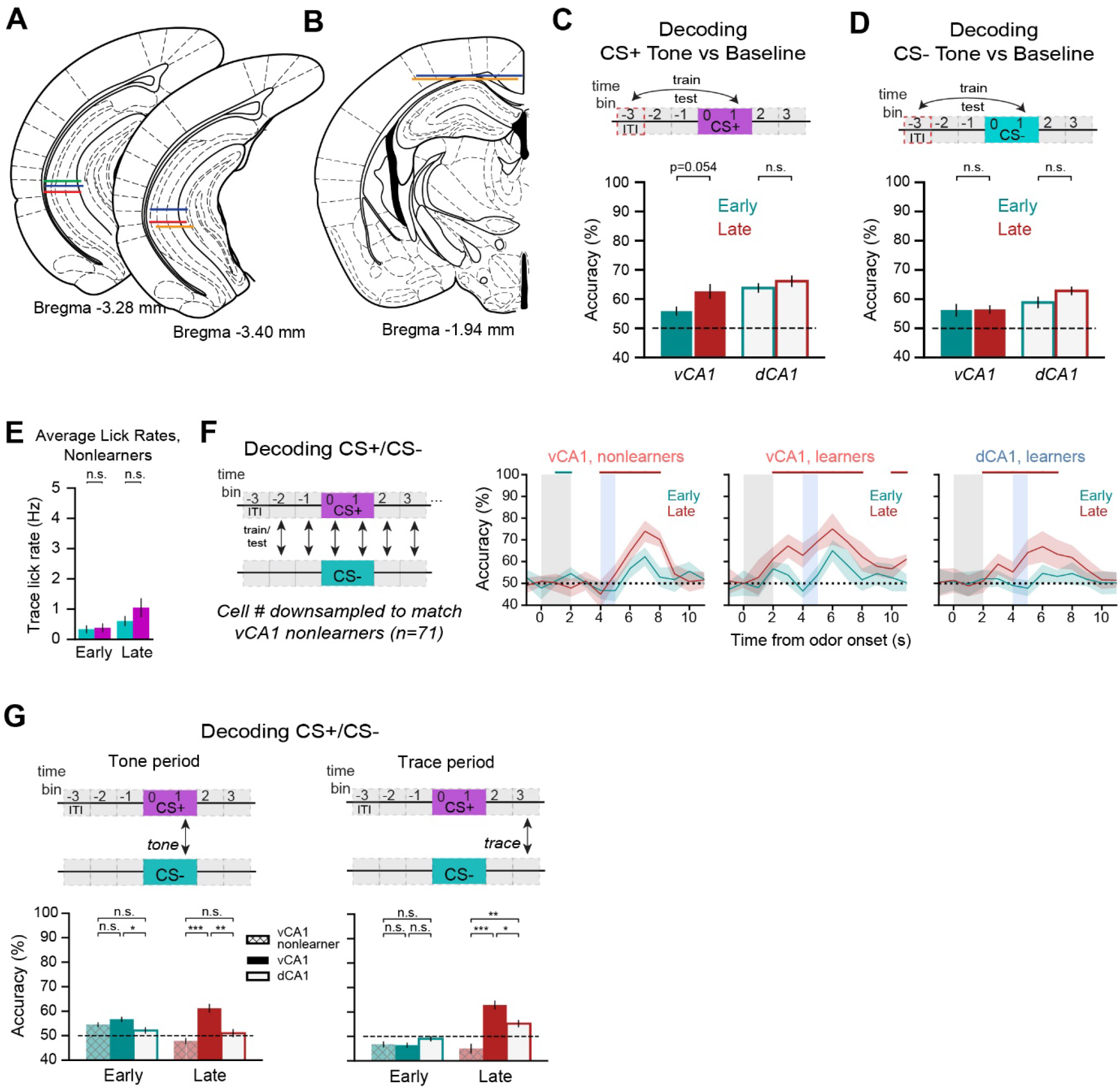
Related to Figure 3. Mice that fail to learn the task do not show representational changes. (A and B) Location of GRIN lens implants in vCA1 (A) and window implants in dCA1 (B) for animals used in the tone associative learning study. Atlas images adapted from (Paxinos and Franklin, 2019) (C and D) Population decoding CS+ (C) or CS-(D) tone vs ITI baseline (E) Average trace-period lick rates for vCA1 animals who failed to learn the discrimination task. (F) Trial-type decoding accuracy (±SD). Because decoding performance is correlated with the number of cells included for analysis (see Figure S2E), we downsampled the number of cells in vCA1 and dCA1 “learners” to each match “nonlearners” (n=71 cells). Color-coded bar above shows periods where the corresponding trial type accuracy is significantly greater than the opposing trial type (p < 0.01, Mann-Whitney U test). Note the absence of odor- and trace-period decoding in nonlearners during the Late session. (G) Trial-type decoding performance during the odor-(left) and trace-period (right) epochs (±SEM, Mann-Whitney U test).

**Figure S4.**
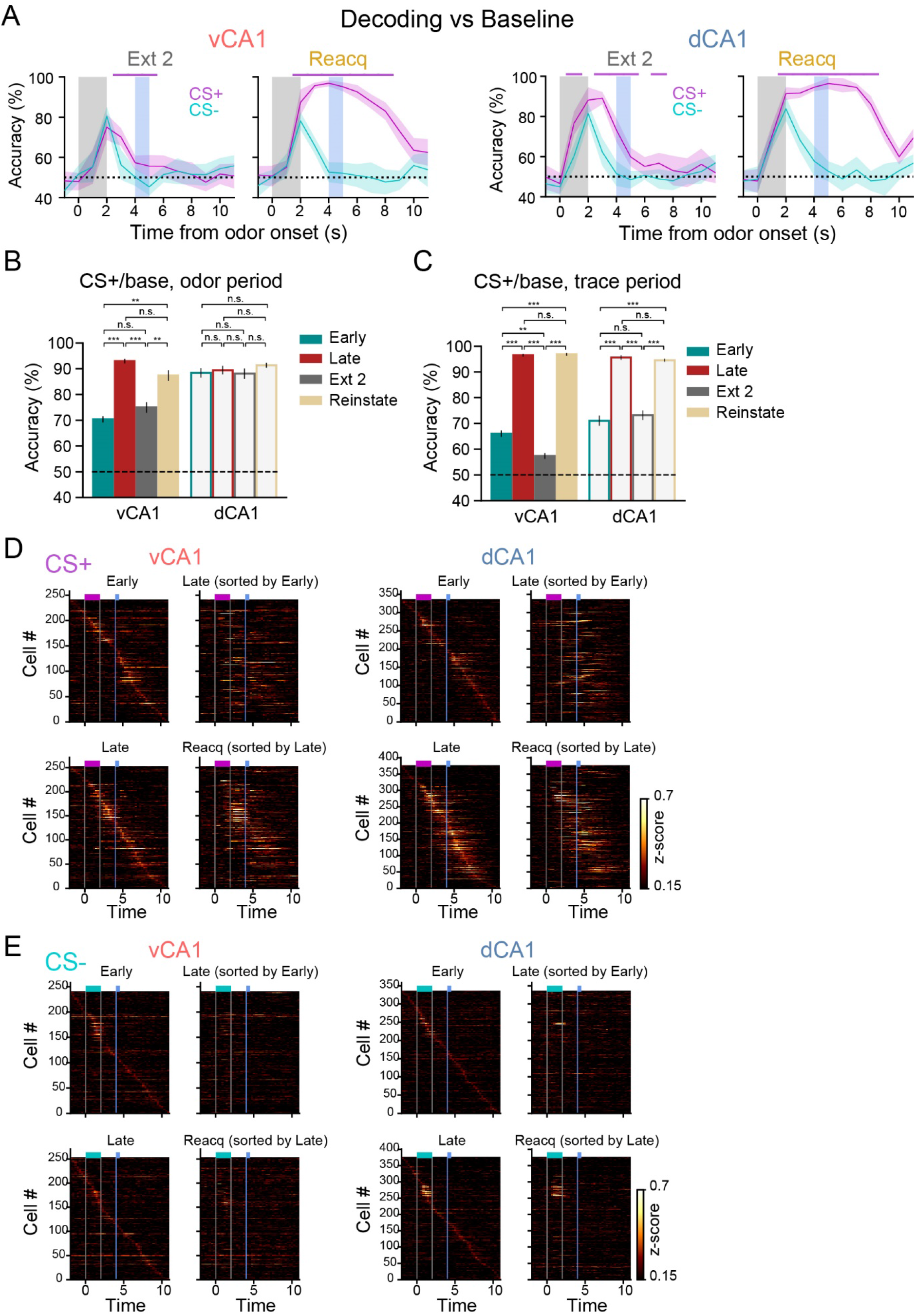
Related to Figure 4. Omitting reward reduces CS+, but not CS-, representations during odor (vCA1) and trace periods (vCA1 and dCA1). (A) Population-activity decoding accuracy for CS+ or CS-trials from baseline (±SD). Color-coded bar above shows periods where the corresponding trial type accuracy is significantly greater than the opposing trial type (p < 0.01, Mann-Whitney U test). (B) CS+ versus baseline odor-period decoding accuracy for each learning phase (±SEM, Mann-Whitney U test). Note that decoding performance is correlated with odor value in vCA1 (left) but not dCA1 (right). (C) Same as (B), but for trace period. Decoding performance is correlated with reward expectation in both vCA1 and dCA1. (D) Activity during CS+ trials for neurons registered across session pairs. For each time bin, activity z-scores for each neuron were averaged across all trials within a session, and neurons were sorted by peak firing rate latency during the indicated session. Note the changing subset of task-responsive cells from Early to Late, and the relative stability following learning (Late to Reacquisition). Also note the few cells responsive to sucrose reward delivery during Early that translocate their firing to the CS and trace periods following CS-US learning. (E) Same as in (D), but for CS-trials.

**Figure S5.**
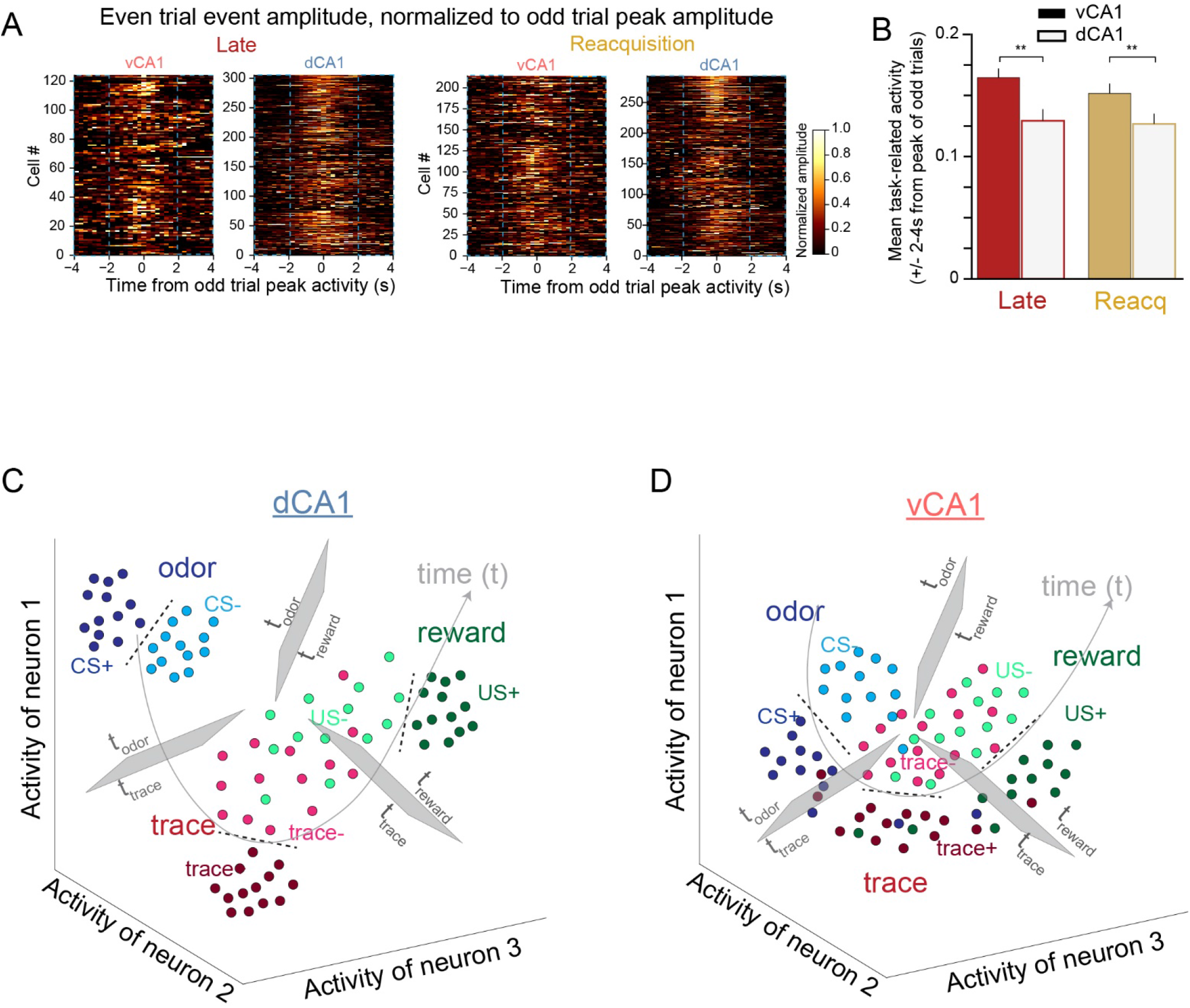
Related to Figure 5. Broad firing in vCA1 and summary schematic. (A) Even trial event amplitude heatmaps for individual neurons whose peak activity during odd trials occurred during the odor or trace periods (see Methods). Data shown are for CS+ trials only. (B) Mean normalized CS+ trial activity collected 2-4 seconds before and after peak of odd trials (corresponding to the data within dashed outlines in (B); ±SEM, Mann-Whitney U test). (C and D) Summary schematic illustrating learned task representations in dCA1 (C) and vCA1 (D). Each dot represents a single-trial population activity vector during odor (blue), trace (red) or US (green) task epochs. In both dCA1 and vCA1, neural representations of CS+ and CS-trials are highly separable during each epoch. Task epochs are also generally separable from one another in both regions, particularly for CS+ trials and comparisons farther apart in time. In dCA1 there is less overlap of representations across epochs than there is in vCA1, owing in part to neural activity with greater temporal specialization in dCA1.

**Figure S6.**
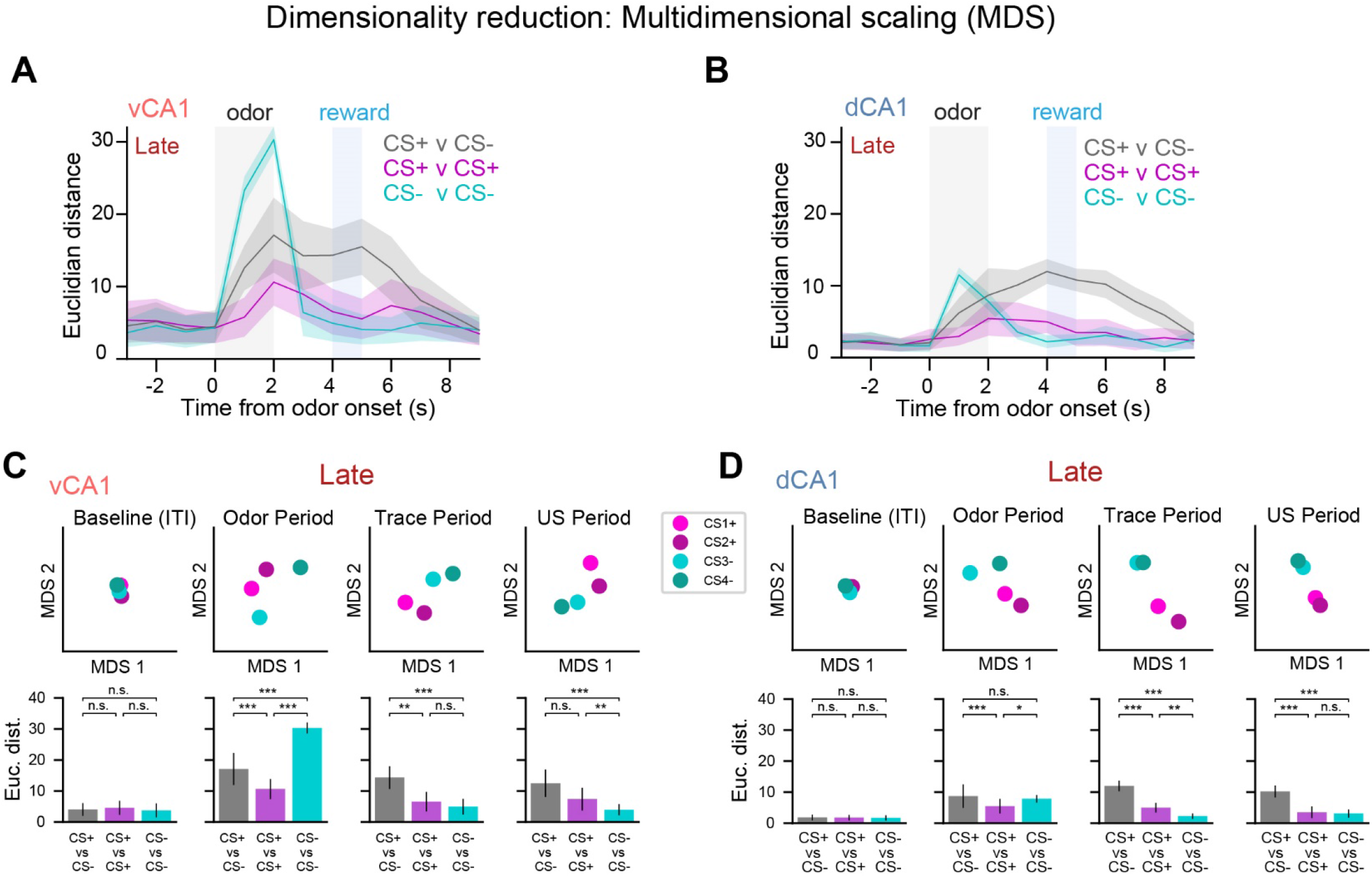
Related to Figure 6. Neural geometry of 4-odor task representations. (A) To analyze the similarity between population activity patterns for each trial type, we projected the hyper-dimensional neural data onto 2-dimensional space via multidimensional scaling (MDS; see Methods). Here, the relationship between activity patterns is represented in geometrical space; the closer two points are in space, the more similar their activity patterns. The graphs show the mean Euclidean distance of 100 MDS runs (±SD) using Late session neural data. Neural representations for each CS diverged from one another upon odor delivery. In contrast, during the trace period each trial type converged closer to their corresponding trial type (e.g., CS1+ with CS2+) but remained distant from opposite trial types (e.g., CS1+ vs CS3-). These data indicate a switch from encoding individual odors to expected outcome (in CS+ trial types) as animals progressed through a trial. (B) Same as (A), but for dCA1. (C) (top) Sample results from a single 2-D MDS run illustrating the relationship of vCA1 population activity for all 4 trial types at different task epochs during the Late session. (bottom) Average Euclidean distance of 100 MDS runs (±SD, Mann-Whitney U test). US Period = 5-6 seconds post-odor onset time bin (1-2 seconds post sucrose delivery). (D) Same as (C), but for dCA1.

**Figure S7.**
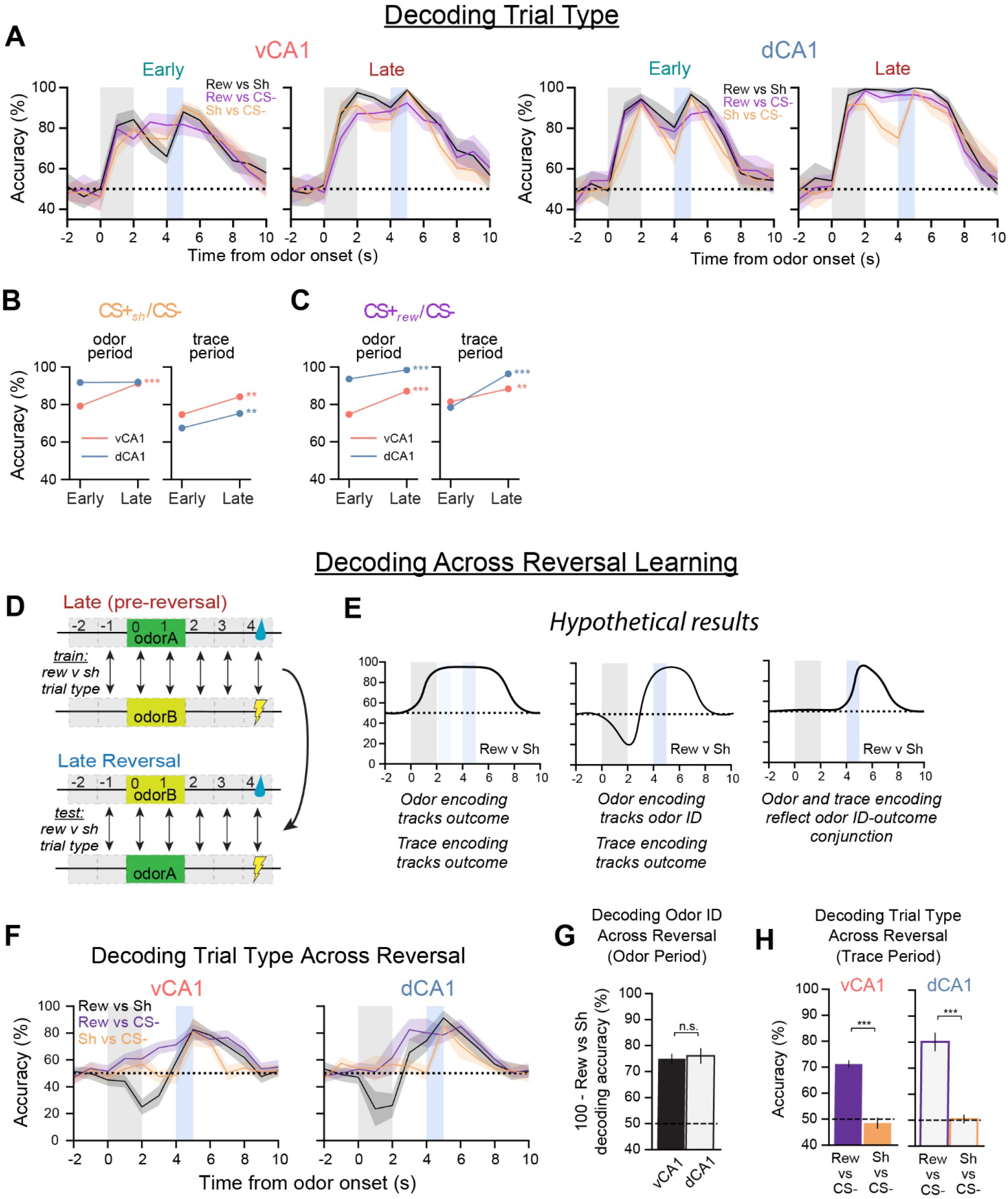
Related to Figure 7. Odor ID and reward expectation representations remain stable across reversal learning, while shock anticipation signals fade. (A) Trial-type decoding accuracy (±SD). (B) Change in odor-period (left) or trace-period (right) decoding accuracies for CS+shock vs CS-trials from Early to Late sessions (±SEM). Statistics compare Early and Late sessions for a specific hippocampal region (Mann-Whitney U test). (C) Same as in (B), but decoding CS+reward from CS-trials. (D) Schematic illustrating trial-type decoding across reversal learning. (E) Hypothetical results for decoding CS+reward from CS+shock trials across reversal learning (for this set of results, stable encoding of US identity across reversal is assumed). Because data classes were labeled with respect to the outcome of a trial, and not the odor identity, stable neural representations of odor identity will manifest as cross-session decoding accuracies that are below chance (middle graph). (F) Actual results for decoding trial type across reversal learning (±SD). The below chance decoding accuracy for CS+reward vs CS+shock during the odor period indicates representations of odor identity dominate the population activity during this time. (G) Across-reversal odor ID decoding accuracy during the odor period (±SEM, Mann-Whitney U test). (H) Across-reversal trial type decoding accuracy during the trace period (±SEM, Mann-Whitney U test).

## Methods

KEY RESOURCES TABLE

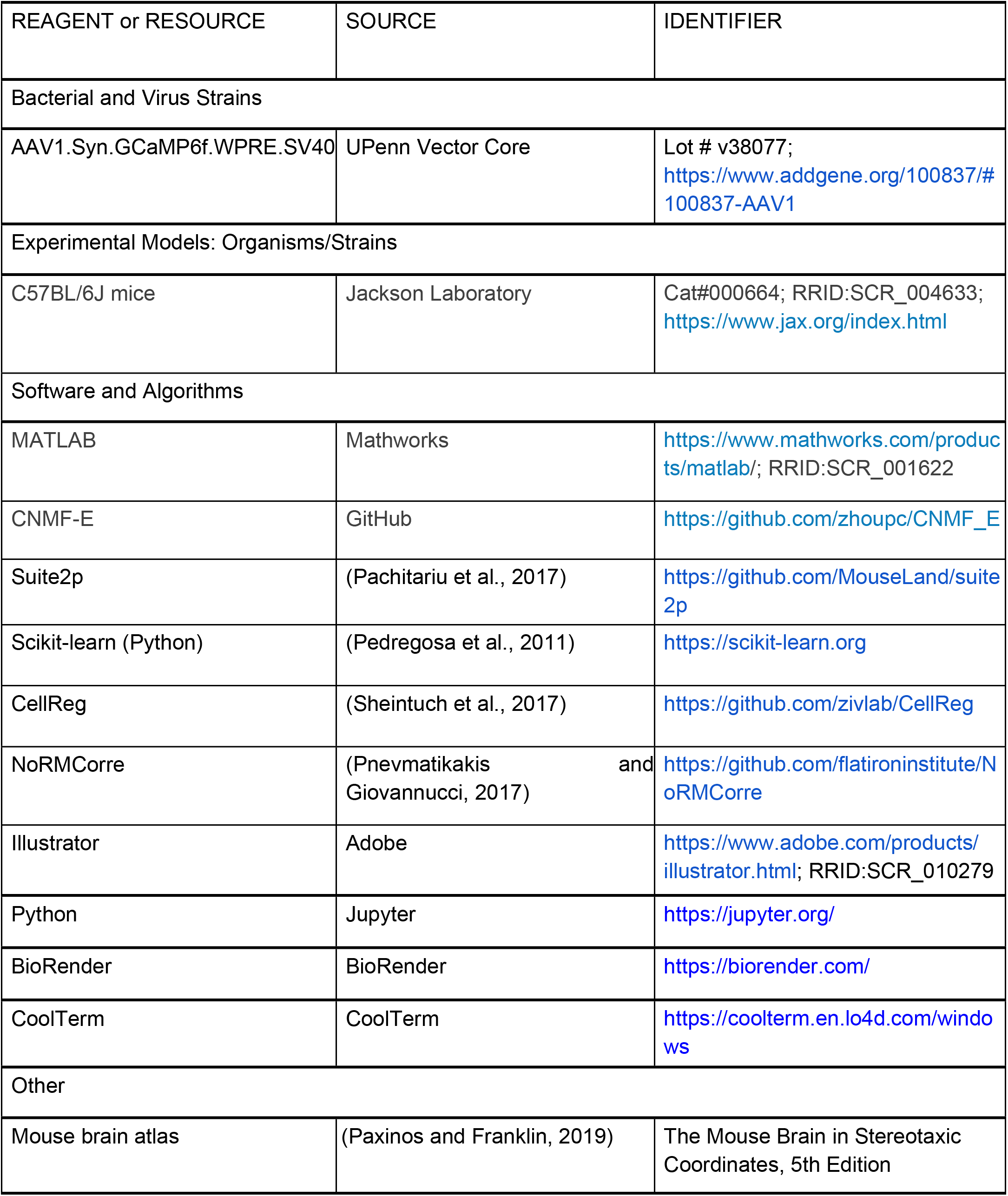

## Contact for reagents and resource sharing

Further information and requests for resources and reagents should be directed to and will be fulfilled by the Lead Contact, Mazen Kheirbek.

## Materials Availability

This study did not generate new unique reagents.

## Data and Code Availability

The datasets and analysis code supporting the current study are available from the lead contact on request.

### Mice

All procedures were conducted in accordance with the NIH Guide for the Care and Use of Laboratory Animals and institutional guidelines. Adult male C57BL/6J mice were supplied by Jackson Laboratory. Mice were kept on a 12-hour light cycle, with experiments conducted during the light portion.

### Surgery

Animals were 11 – 15 weeks postnatal at time of surgery. Mice were anesthetized with 1.5% isoflurane with an O2 flow rate of 1 L / min, and head-fixed in a stereotactic frame (David Kopf, Tujunga, CA). Eyes were lubricated with an ophthalmic ointment, and body temperature was maintained at 37°C with a warm water recirculator (Stryker, Kalamazoo, MI). The fur was shaved and incision site sterilized prior to beginning surgical procedures. Lidocaine, meloxicam and slow-release buprenorphine were provided for analgesia.

GCaMP6f virus injection and GRIN lens implantation were conducted using methods previously described (Jimenez et al., 2018). Briefly, a craniotomy was made over the lens implantation site and dura was removed from the brain surface and cleaned with sterile saline and absorptive spears (Fine Science Tools, Foster City, CA). A nanoject syringe (Drummond Scientific, Broomall, PA) was used to deliver GCaMP6f to vCA1 or dCA1 (left hemisphere for both). vCA1 coordinates were −3.16 A/P and −3.25 M/L. 150nl of virus was injected at each depth of −3.85, −3.55 and −3.3 (450nl total volume) with respect to bottom of skull at the medial edge of the craniotomy. dCA1 coordinates were −2 A/P, −1.65 M/L and −2.1 A/P, −1.45 M/L at depths −1.5, −1.25 D/V with respect to bregma. The needle was held in place for > 5 minutes prior to moving to the next D/V coordinate, and remained in place for 10 minutes following the final injection before slowly removing from the brain. AAV1-SYN-GCaMP6f-WPRE-Sv40 (titer: 1.97E+13) was supplied from University of Pennsylvania viral vector core and diluted 1:3 in 1x sterile PBS before injections. For dCA1, prior to virus injection the overlying cortex was slowly aspirated until axonal fibers of the external capsule/alveus were visualized. Following virus injection, a 0.6mm (vCA1) or 1.0mm (dCA1) diameter GRIN lens (Inscopix, Palo Alto, CA) was slowly lowered in 0.1 mm D/V steps and then fixed to the skull with Metabond adhesive cement (Parkell, Edgewood, NY). vCA1 lens coordinates were −3.16 A/P, −3.5 M/L and −3.5 D/V (from bottom of skull at craniotomy; Figures S1A and S3A). dCA1 lens coordinates were −2.05 A/P, −1.5 M/L, −0.95 D/V (from bregma; Figure S1B). A custom-made titanium headbar was then attached to the skull using dental cement (Dentsply Sinora, Philadelphia, PA). A baseplate and cover (Inscopix, Palo Alto, CA) was also cemented on to protect the lens.

For dCA1 animals in the tone discrimination paradigm, a 3mm craniotomy was made, and the overlying cortex was aspirated until axonal fibers of the external capsule/alveus were visualized. The aspiration site was continuously irrigated with cold, sterile saline. Viral injections (120 nl per site) were performed at the same sites as above. A custom made dCA1 imaging window was implanted, which consisted of a 3mm round coverslip, #0 thickness (Warner Instruments, Hamden, CT) attached with optical adhesive (#81, Norland Products, Cranbury, NJ) to a metal cannula containing 1/8” outer diameter and 1/16” in length (McMaster-Carr, Santa Fe Springs, CA). This window was carefully lowered into place, until it rested on top of the exposed tissue (Figure S3B). The cannula was then cemented into place with Metabond adhesive, and a custom titanium headbar was cemented in place.

### Verification of imaging sites and histological analysis

Dorsal and ventral CA1 imaging sites were verified in each animal included in final analysis (Figures S1 and S3). After all imaging sessions were completed, mice were injected with a lethal dose 2:1 ketamine/xylazine solution intraperitoneally. While the heart was still beating, mice were perfused transcardially using 4% PFA solution. Brains were extracted and placed in 4% PFA solution for 2-3 days to allow further fixation. After saturating with a 30% sucrose solution, coronal slices of 50-micron width were collected using a Leica SM2000 microtome. Slices were collected in 1x PBS solution and mounted onto glass slides, coverslipped with Fluoromount G with DAPI (Southern Biotech, Birmingham, AL).

### Behavioral training

Four-to-six weeks following surgery, animals were handled and habituated to the experimenter, training environment and head fixation for one week. Following habituation, animals were water restricted to ∼85-90% ad lib weight and underwent a 2-3 day pretraining period designed to introduce the sucrose delivery apparatus, with free sucrose rewards (∼2 µl each) intermittently delivered upon licking (up to 80 rewards in a 20 min session). Sucrose rewards (10% sucrose, 0.03% NaCl in water) were delivered via a solenoid-gated gravity feed. Contact with a lick spout positioned in front of the mouth was measured using a capacitive touch MPR121 sensor (SparkFun, Boulder, CO). Stimulus delivery and sensor reading was controlled by an Arduino Mega with custom circuit boards (adapted from OpenMaze.org) and recorded via CoolTerm software. Once animals displayed consistent and motivated licking (80 rewards collected in a single session), lick training was complete and the pretraining odor exposure session was initiated the following day. Throughout training, animals were water restricted to ∼85-90% ad lib weight. All training paradigms consisted of one training session/day, occurring at roughly the same time each day. Learning of the discrimination tasks was assessed using lick discriminability (*d’*) for each session, which compares the rate of anticipatory licks during the trace period of CS+ trials with CS-trials:

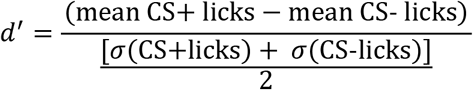

Learning was determined as a *dd*’ score > 1.5 for a session, with all Late session mice meeting this criterion.

#### Pretraining odor exposure

One day prior to conditioning, animals underwent a single session where they were passively exposed to neutral odors (benzaldehyde, eugenol) that would subsequently serve as CS+ and CS-odors in the 2-odor paradigm. Each session consisted of 30 trials (15 of each odor) of 2 second odor presentations. There was no lick spout present during this session. The inter-trial interval between subsequent odor deliveries was chosen as a random sample from a uniform distribution between 17.5 and 27 seconds. Odors were delivered via a custom-made olfactometer equipped with a mass flow controller (Alicat Scientific, Tucson, AZ) that maintained air flow at 2 liters per minute and prevented momentary pressure changes from solenoid valve switches (Clippard, Cincinnati, OH) upstream of the controller. Odors were delivered to mice via a customized nose cone, which contained an outlet where a gentle vacuum was applied to evacuate residual odor. Additionally, an ongoing charcoal filter vacuum system (Hydrobuilders Inc.) was used to evacuate any residual odors.

#### 2-odor paradigm

Each associative learning session consisted of 120 trials (60 CS+ and 60 CS-, pseudorandomly presented). Two neutral odors served as CS+ and CS-cues (benzaldehyde or eugenol, 2s) with cue contingencies counterbalanced across mice. Presentation of the CS+ cue was followed by a 2s trace period and subsequent reward delivery (∼2 µl). No reward was available following the presentation of the CS-cue. Animals were not punished for off-target licking. The inter-trial interval between subsequent cues was chosen as a random sample from a uniform distribution between 17.5 and 27 seconds.

This task structure was administered over a period of ∼7 days, in which day 1 and 4 were termed “Early” and “Late” learning, respectively. If an animal did not meet the learning criterion (d’ > 1.5) on day 4 (n=3 animals), training continued until this criterion was met. The two days following the Late session extinction sessions, labeled as “Ext1” and “Ext2”, respectively, in which the odor-reward association was extinguished by removing the sucrose reward for CS+ trials (the lick spout remained in place). Ext2 was followed by a one-day reacquisition session, labeled as “Reacquisition”, in which the sucrose reward was reintroduced for CS+ trials. A total of 11 vCA1 and 5 dCA1 animals were included in the data set.

#### 2-tone paradigm

Each associative learning session consisted of 160 trials (80 CS+ and 80 CS-, pseudorandomly presented). Two auditory tones served as CS+ and CS-cues (2.5 kHz and 13kHz pulsing tones, 2s, 70 dBs) with cue contingencies counterbalanced across mice. Presentation of the CS+ cue was followed by a 2s trace period, then a 2s reward window which required a lick for sucrose reward delivery (∼2 µl, maximum one reward per trial). No reward was available following the presentation of the CS-cue. Animals were not punished for off-target licking. The inter-trial interval between subsequent cues was chosen as a random sample from a uniform distribution between 17.5 and 27 seconds. A total of 7 vCA1 and 2 dCA1 animals were included for analysis (a separate cohort of mice than that used for the odor-based experiments). In a subset of animals (n= 4 vCA1 and n = 2 dCA1), multiple z-planes were imaged across sessions. Imaging planes were separated by > 60 µm to ensure there was no overlap of cells present across different z-planes.

#### 3-odor paradigm

Following completion of 2-odor training, mice underwent 3-odor conditioning. Each session consisted of 120 trials (40 CS+rew. 40 CS+shock, 40 CS-, pseudorandomly presented). Three new neutral odors served as cues (o-toluidine, 2-heptanone, or +carvone; 2s). Presentation of the CS+ cue was followed by a 2s trace period and subsequent delivery of US (reward US = ∼2 µl 10% sucrose solution; shock US = 0.125mA amplitude, 250ms duration). No US was presented following the presentation of the CS-cue. Animals were not punished for off-target licking. For reversal learning, CS+reward and CS+shock odor cues were switched, while the CS-odor remained the same. Shocks were delivered via a custom-made tail cuff driven by a precision animal shocker (Coulbourn Instruments, Holliston, MA). The inter-trial interval between subsequent cues was chosen as a random sample from a uniform distribution between 17.5 and 27 seconds. A total of 11 vCA1 and 3 dCA1 animals were included in the data set.

#### 4-odor paradigm

Following 3-odor training, mice underwent 4-odor conditioning. Each session consisted of 120 trials (30 of each trial type, pseudorandomly presented). Four neutral odors served as cues (methylbutyrate, isoamyl acetate, eugenol, eucalyptol), one of which (eugenol) had been experienced previously in the 2-odor task (due to a lack of available neutral odors) and was paired with the same outcome as previously experienced. Presentation of the CS+ cue was followed by a 2s trace period and subsequent delivery of 10% sucrose solution (∼2 µl). No US was presented following the presentation of the CS-cue. Animals were not punished for off-target licking. The inter-trial interval between subsequent cues was chosen as a random sample from a uniform distribution between 17 and 23 seconds. A total of 7 vCA1 and 3 dCA1 animals were included in the data set.

#### 2-photon imaging

Genetically encoded calcium imaging of GCaMP6f was used to assess the functional activity of individual neurons. Images were captured using an Ultima IV laser scanning microscope (Bruker Nano, Middleton, WI) equipped with resonant scanning mirrors and high-speed scan electronic controller, dual GaAsP PMTs (Hamamatsu model 7422PA-40), and motorized z focus (100 nm step size). GCaMP signal was filtered through an ET-GFP (FITC/CY2) filter set. Laser signal was provided by a MaiTai DeepSee mode-locked Ti:Sapphire laser source (Spectra-Physics, Irvine, CA) providing > 150kW max output at 920 nm. Acquisition speed was 30Hz for 512 × 512 pixel images. Images were averaged both online and offline, yielding a final frame rate of 3.75Hz.

Prior to each conditioning session, the imaging field of view (FOV) was determined, and imaging was conducted at that FOV for the entire session. For animals with multiple FOVs across sessions, each FOV was separated by > 60 µm in the z-dimension (dorsal-ventral) to ensure no overlap of cells across different FOVs. To facilitate re-identification of a specific FOV across sessions, the top of the GRIN lens served as a reference z-plane. Optimal laser power was determined for each FOV based on GCaMP expression level and was kept constant across sessions for a specific FOV. For each trial, imaging began 8 sec prior to cue onset and was terminated 11 sec afterward (19 sec total).

### Signal extraction and cross-session registration

Videos were motion corrected offline using non-rigid motion correction based on template matching (NoRMCorre (Pnevmatikakis and Giovannucci, 2017) or Suite2p (Pachitariu et al., 2017)). Cell segmentation and calcium transient time series data were extracted using Constrained Non-negative Matrix Factorization for microEndoscopic data (CNMF-E), a semi-automated algorithm optimized for GRIN lens Ca2+ imaging to denoise, deconvolve and demix calcium imaging data (Zhou et al., 2018). Putative neurons were manually inspected for appropriate spatial properties and Ca2+ dynamics, and were visually checked against the corresponding motion corrected video in ImageJ. Ca2+ transient events were extracted using the OASIS algorithm (Friedrich et al., 2017) embedded within CNMFe. We used these inferred calcium events for all analyses, unless otherwise noted.

Registration of cells across sessions imaged at the same FOV used probabilistic modeling of similarities between cell pairs across sessions (CellReg, (Sheintuch et al., 2017)). Briefly, spatial footprint maps were generated for each session by projecting the spatial filter of each cell onto a single image. Spatial footprint images from sessions imaged at the same FOV were then aligned. The distribution of similarities between pairs of neighboring cells were subsequently modeled via centroid distance to obtain an estimation for their probability of being the same cell (P_same_). Cells were then registered across sessions via a clustering procedure that utilizes the previously obtained probabilities, with a probability threshold of 0.8. The average P_same_ value for registered cells was 0.95. All putative matches were visually inspected.

### Data analysis

For statistical analyses and figures, calcium event activity was separated into 1-second bins and average activity during each bin was used. When reporting specific epochs of task results, “odor period” constituted the final 1-second bin of odor delivery (1-sec to 2-sec post odor onset), while “trace period” constituted the final 1-second bin of the trace period prior to reward delivery (1-sec to 2-sec post odor offset), unless otherwise noted. These time bins were chosen to ensure odor was being experienced throughout the entire odor bin and to minimize any residual odor effects during trace period analysis.

#### Population decoding

A linear decoder was used to discriminate activity patterns into two discrete categories (Bishop, 2006):

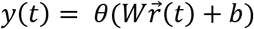

where *y* is the predicted label of the population activity pattern 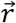 recorded at time 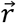 and takes two values corresponding to two classes of patterns to decode (for example, two odor identities), *W* is the vector of weights assigned to each cell, and *b* is a bias term constant. Decoding parameters were attained via a supervised learning protocol with labeled data and used a support-vector machine (SVM) with a linear kernel (python/scikit/linearSVC). Results are reported as the generalized performance of the decoder using cross-validation, a standard machine learning procedure to avoid data overfitting. When multiple categories were involved, e.g., more than two trial types, multiple linear decoders were trained on pairs of discrete categories combined using majority-based error-correction codes.

We defined the patterns of calcium activity by computing the mean event rates during one-second time bins. Pseudo-population recordings were generated by combining cells across multiple animals/FOVs. For decoding, one-half of trials were randomly selected from each class and pseudo-population activity from these trials was used to train the decoder, while the remaining held-out half was used to evaluate the decoder’s generalization performance. When comparing decoding accuracy between neural populations of different size, we trained our decoder on a random subsample of cells from the more numerous population equal to that of the smaller population. We repeated the operation 100 times and then combined the cross-validated decoding accuracies of all random choices together to get a single sample of decoding accuracies. We repeated the procedure 10 times to perform statistical comparisons across groups and against chance performance. A two-sided Mann-Whitney U Test was used to compare decoding accuracies between groups, and Bonferroni correction used for multiple comparisons.

For decoding against baseline, we used population activity during the 1-second time bin that began three seconds prior to CS onset as baseline data. Cross-time-bin and cross-session decoding followed the same procedure as within-session decoding. In the case of cross-session decoding, only cells registered across the compared sessions were included.

For decoding distant time bins in our cross-time-bin decoding analysis (Figures 6C, 7H and 7J), we took the average of all decoding runs for each cross-time-bin comparison that was separated by 3 or more bins. Further, we only included comparisons where both train and test data occurred at least 1 second after odor onset (that is, pre-trial data was excluded).

#### Multidimensional Scaling (MDS)

We performed 2-dimensional MDS scaling of event data using python/scikit/MDS. As with decoding, we combined all cells recorded from a particular region (e.g., vCA1) across all mice into one pseudopopulation. For each trial type, 100 trials were randomly selected for analysis, and MDS was performed. The Euclidean distance was taken between each trial type, and this process was repeated 100 times. Bar charts of Euclidean distance show the mean ± SD of all runs.

#### Single-cell responsivity

Data used for heatmaps of calcium-traces or inferred events were not binned. For each cell, z-scores were computed over the entire dataset for a specific condition (e.g., CS+ trials). To identify cells whose activity was modulated during specific epochs (e.g., CS+ period, trace period, etc), for each trial containing the specified epoch the average event magnitude during the 1s epoch was compared to the average event magnitude during a 1s baseline period immediately prior to cue onset for that trial. P-values were determined using a two-tailed Mann Whitney U test and the False Discovery Rate (FDR) was applied to correct for multiple comparisons. Cells with an adjusted p-value < 0.05 were classified as responsive. Fisher’s exact test was used to compare whether the proportion of selective cells for a specific epoch (eg, CS+ Early vs CS+ Late) significantly differed (p < 0.05).

To compare the persistence of CS+-trial-related activity in vCA1 vs dCA1 neurons, we first parsed CS+ trial data into odd or even trials and averaged activity for each cell across these trials. We then extracted cells whose peak activity during odd trials occurred between odor onset and US onset. Average activity for these cells during even trials was then collected +/-4 seconds around the time point of odd-trial peak activity, normalized to the amplitude of odd-trial peak activity, and plotted (Figure S5A).

#### Lick Correlation

We determined whether activity of dCA1 or vCA1 neurons were correlated with licking. We regressed the lick rates across the session against calcium events. We fit a linear regression model to predict lick rates and used the explained variance (R^2^) as a measure of goodness of fit to compare the results across animals and days. We divided each analyzed session in 10 time-contiguous blocks and computed the generalization performance of the model with 10-fold cross-validation over these blocks to avoid overfitting. Regression was performed with regular linear regression with Lasso, and verified that the results are not qualitatively different.

#### Aha Analysis

We identified the first moment of distinguished licking behavior between CS+ and CS-trials by locating in all mice in the 2-odor paradigm an “aha” moment. This was calculated by averaging across every 4 trials the cumulative CS+ and CS-lick rates, taking the slope of the difference in cumulative licking between these bins, and checking if 1) the difference exceeded the previous bin’s slope >= 1 standard deviation of the difference line up to that bin, and 2) the slope increase exceeded 1/3 of the difference between the previous set of trials. The averaging and thresholding with an increased slope relative to previous trials limited detection of instances where a short sequence of successfully discriminated trials were followed by a return to incorrect lickings, which would not represent a true aha moment. For potential aha moments detected on the first day of learning, we set a threshold of a minimum of 80 licks so that only mice who demonstrated lick rates similar to or above the baseline we required during lick training could be considered to have learned. All aha moments detected by this method were cross-checked with examining the raw licking data to ensure accuracy. Aha moments across mice spanned the first two days of learning, with 62% of mice reaching an aha moment on the first day or learning, and all mice reaching an aha moment by the end of the second day.

For aha population decoding analysis, we used 30 trials before or after the aha moment. For mice where the aha moment was < 30 trials from the end of the first or beginning of the second day of learning, trials from both days were included in order to reach the full 30 trials, and only cells registered across both sessions were included.

#### Active time bins analysis

For this analysis, time series data were binned into 0.25 sec time bins to provide greater resolution. Because the Pre session only contained 15 trials of each trial type, only the first 15 trials of each trial type were included for all sessions examined. For each trial (defined as 0 - 8 seconds post odor onset) and each time bin within that trial (32 time bins total) we examined whether an event was present. The data presented show the number of time bins where an event was detected for at least one trial. For example, a cell that fired during time bin 5, and only time bin 5, on every trial would produce a score of 1. A cell firing on time bin 5 and 10 on trial 1, bins 7 and 10 on trial 7, and no firing on all other trials would produce a score of 3. A maximum score of 32 indicates that each of the 32 individual time bins registered an event on at least one of the 15 trials. This analysis was repeated for each cell. To compare whether individual cells changed with learning, we only included cross-registered cells.

For all figures: * p< 0.05, ** p < 0.01, *** p < 0.001. See Table S1 for all statistical analysis details.

