## Supplemental Table S1 for "Population dynamics underlying associative learning in the dorsal and ventral hippocampus"

| Figure | Variable | Unit of Comparison | n | Test | Results |
|---|---|---|---|---|---|
| Fig. 1G | decoding accuracy vs chance | 10 decoding iterations for each region | pseudopopulation (see methods) of 454 cells (n-matched in vCA1 and dCA1) from 11 vCA1 and 5 dCA1 mice | Wilcoxon | color-coded bars above graph show time bins where p < 0.01 |
| Fig. 1H | vCA1 vs dCA1 odor1/baseline decoding accuracies | 10 decoding iterations for each region | n-matched pseudopopulation of 454 cells from 11 vCA1 and 5 dCA1 mice | Mann-Whitney U | U = 0, p = 0.00017, effect-size (r) = 0.85 |
| Fig. 1H | vCA1 vs dCA1 odor2/baseline decoding accuracies | 10 decoding iterations for each region | n-matched pseudopopulation of 454 cells from 11 vCA1 and 5 dCA1 mice | Mann-Whitney U | U = 12.0, p = 0.0046, effect-size (r) = 0.64 |
| Fig. 1J | odor1/odor2 decoding accuracy, vCA1 vs dCA1 | 10 decoding iterations for each region | n-matched pseudopopulation of 454 cells from 11 vCA1 and 5 dCA1 mice | Mann-Whitney U | colored-coded bar above graph shows time bins where p < 0.01 |
| Fig. 2D | mean lick rate (Hz) | Trial type (Early session) | 15 mice (11 vCA1 and 4 dCA1 mice) | Mann-Whitney U | U =75.5, p =0.13, effect-size (r) = 0.28 |
| Fig. 2D | mean lick rate (Hz) | Trial type (Late session) | 15 mice (11 vCA1 and 4 dCA1 mice) | Mann-Whitney U | U =0, p < 0.0001, effect-size (r) = 0.85 |
| Fig. 2G | CS+/baseline vs CS-/baseline decoding accuracies, vCA1 or dCA1 | 10 decoding iterations for each trial type | n-matched pseudopopulation of 454 cells from 11 vCA1 or 4 dCA1 mice | Mann-Whitney U | color-coded bars above graph show time bins where p < 0.01 |
| Fig. 2H | vCA1 vs dCA1 CS+/baseline decoding accuracies during odor period (Pre session) | 10 decoding iterations for each region | n-matched pseudopopulation of 454 cells from 11 vCA1 and 4 dCA1 mice | Mann-Whitney U | U =0, p = 0.00017, effect-size (r) = 0.85 |
| Fig. 2H | vCA1 vs dCA1 CS+/baseline decoding accuracies during odor period (Late session) | 10 decoding iterations for each region | n-matched pseudopopulation of 454 cells from 11 vCA1 and 5 dCA1 mice | Mann-Whitney U | U =69.5, p = 0.15, effect-size (r) = 0.33 |
| Fig. 2H | vCA1 vs dCA1 CS+/baseline decoding accuracies during trace period (Pre session) | 10 decoding iterations for each region | n-matched pseudopopulation of 454 cells from 11 vCA1 and 4 dCA1 mice | Mann-Whitney U | U =25.5, p = 0.069, effect-size (r) = 0.41 |
| Fig. 2H | vCA1 vs dCA1 CS+/baseline decoding accuracies during trace period (Late session) | 10 decoding iterations for each region | n-matched pseudopopulation of 454 cells from 11 vCA1 and 5 dCA1 mice | Mann-Whitney U | U =56, p = 0.68, effect-size (r) = 0.10 |
| Fig. 2J | vCA1 vs dCA1 CS+/CS- decoding accuracies during odor period (Pre session) | 10 decoding iterations for each region | n-matched pseudopopulation of 454 cells from 11 vCA1 and 4 dCA1 mice | Mann-Whitney U | U = 0, p = 0.00018, effect-size (r) = 0.85 |
| Fig. 2J | vCA1 vs dCA1 CS+/CS- decoding accuracies during odor period (Late session) | 10 decoding iterations for each region | n-matched pseudopopulation of 454 cells from 11 vCA1 and 5 dCA1 mice | Mann-Whitney U | U = 44, p = 0.68, effect-size (r) = 0.10 |
| Fig. 2J | vCA1 vs dCA1 CS+/CS- decoding accuracies during trace period (Pre session) | 10 decoding iterations for each region | n-matched pseudopopulation of 454 cells from 11 vCA1 and 4 dCA1 mice | Mann-Whitney U | U = 12, p = 0.005, effect-size (r) = 0.64 |
| Fig. 2J | vCA1 vs dCA1 CS+/CS- decoding accuracies during trace period (Late session) | 10 decoding iterations for each region | n-matched pseudopopulation of 454 cells from 11 vCA1 and 5 dCA1 mice | Mann-Whitney U | U = 32, p = 0.18, effect-size (r) = 0.30 |
| Fig. 2L | within-session Early vs within-session Late CS+/CS- decoding accuracies during odor period (vCA1) | 10 decoding iterations for each | n-matched pseudopopulation of 241 cells from 11 vCA1 and 4 dCA1 mice | Mann-Whitney U, Bonferroni correction for n=2 | U = 0, p < 0.001, effect-size (r) = 0.85 |
| Fig. 2L | within-session Early vs across-session Early/Late CS+/CS- decoding accuracies during odor period (vCA1) | 10 decoding iterations for each | n-matched pseudopopulation of 241 cells from 11 vCA1 and 4 dCA1 mice | Mann-Whitney U, Bonferroni correction for n=2 | U = 87, p = 0.,011 effect-size (r) = 0.63 |
| Fig. 2L | within-session Late vs across-session Early/Late CS+/CS- decoding accuracies during odor period (vCA1) | 10 decoding iterations for each | n-matched pseudopopulation of 241 cells from 11 vCA1 and 4 dCA1 mice | Mann-Whitney U, Bonferroni correction for n=2 | U = 0, p < 0.001, effect-size (r) = 0.85 |
| Fig. 2L | within-session Early vs within-session Late CS+/CS- decoding accuracies during odor period (dCA1) | 10 decoding iterations for each | n-matched pseudopopulation of 241 cells from 11 vCA1 and 4 dCA1 mice | Mann-Whitney U, Bonferroni correction for n=2 | U = 56, p = 1, effect-size (r) = 0.1 |
| Fig. 2L | within-session Early vs across-session Early/Late CS+/CS- decoding accuracies during odor period (dCA1) | 10 decoding iterations for each | n-matched pseudopopulation of 241 cells from 11 vCA1 and 4 dCA1 mice | Mann-Whitney U, Bonferroni correction for n=2 | U = 99, p < 0.001, effect-size (r) = 0.83 |
| Fig. 2L | within-session Late vs across-session Early/Late CS+/CS- decoding accuracies during odor period (dCA1) | 10 decoding iterations for each | n-matched pseudopopulation of 241 cells from 11 vCA1 and 4 dCA1 mice | Mann-Whitney U, Bonferroni correction for n=2 | U = 0, p < 0.001, effect-size (r) = 0.85 |
| Fig. 2L | within-session Early vs within-session Late CS+/CS- decoding accuracies during trace period (vCA1) | 10 decoding iterations for each | n-matched pseudopopulation of 241 cells from 11 vCA1 and 4 dCA1 mice | Mann-Whitney U, Bonferroni correction for n=2 | U = 2, p < 0.001, effect-size (r) = 0.81 |
| Fig. 2L | within-session Early vs across-session Early/Late CS+/CS- decoding accuracies during trace period (vCA1) | 10 decoding iterations for each | n-matched pseudopopulation of 241 cells from 11 vCA1 and 4 dCA1 mice | Mann-Whitney U, Bonferroni correction for n=2 | U = 75, p = 0.13, effect-size (r) = 0.42 |
| Fig. 2L | within-session Late vs across-session Early/Late CS+/CS- decoding accuracies during trace period (vCA1) | 10 decoding iterations for each | n-matched pseudopopulation of 241 cells from 11 vCA1 and 4 dCA1 mice | Mann-Whitney U, Bonferroni correction for n=2 | U = 100, p < 0.001, effect-size (r) = 0.85 |
| Fig. 2L | within-session Early vs within-session Late CS+/CS- decoding accuracies during trace period (dCA1) | 10 decoding iterations for each | n-matched pseudopopulation of 241 cells from 11 vCA1 and 4 dCA1 mice | Mann-Whitney U, Bonferroni correction for n=2 | U = 0, p < 0.001, effect-size (r) = 0.85 |
| Fig. 2L | within-session Early vs across-session Early/Late CS+/CS- decoding accuracies during trace period (dCA1) | 10 decoding iterations for each | n-matched pseudopopulation of 241 cells from 11 vCA1 and 4 dCA1 mice | Mann-Whitney U, Bonferroni correction for n=2 | U = 64, p = 0.61, effect-size (r) = 0.24 |
| Fig. 2L | within-session Late vs across-session Early/Late CS+/CS- decoding accuracies during trace period (dCA1) | 10 decoding iterations for each | n-matched pseudopopulation of 241 cells from 11 vCA1 and 4 dCA1 mice | Mann-Whitney U, Bonferroni correction for n=2 | U = 100, p < 0.001, effect-size (r) = 0.85 |

| Figure | Variable | Unit of Comparison | n | Test | Results |
|---|---|---|---|---|---|
| Fig. 3C | mean lick rate (Hz) | Trial type (Early session) | 6 mice (4 vCA1, 2 dCA1) | Mann-Whitney U | U =82, p =0.21, effect-size (r) = 0.22 |
| Fig. 3C | mean lick rate (Hz) | Trial type (Late session) | 6 mice (4 vCA1, 2 dCA1) | Mann-Whitney U | U =0, p < 0.0001, effect-size (r) = 0.85 |
| Fig. 3D | decoding accuracy vs chance, vCA1 vs dCA1 | 10 decoding iterations for each region | n-matched pseudopopulation of 537 cells from 4 vCA1 and 2 dCA1 mice | Wilcoxon | color-coded bars above graph show time bins where p < 0.01 |
| Fig. 3E | vCA1 vs dCA1 CS+/baseline decoding accuracies during tone period (Early session) | 10 decoding iterations for each region | n-matched pseudopopulation of 537 cells from 4 vCA1 and 2 dCA1 mice | Mann-Whitney U | U = 15, p = 0.009, effect-size (r) = 0.59 |
| Fig. 3E | vCA1 vs dCA1 CS+/baseline decoding accuracies during tone period (Late session) | 10 decoding iterations for each region | n-matched pseudopopulation of 537 cells from 4 vCA1 and 2 dCA1 mice | Mann-Whitney U | U = 38, p = 0.38, effect-size (r) = 0.2 |
| Fig. 3E | vCA1 vs dCA1 CS+/baseline decoding accuracies during trace period (Early session) | 10 decoding iterations for each region | n-matched pseudopopulation of 537 cells from 4 vCA1 and 2 dCA1 mice | Mann-Whitney U | U = 89, p = 0.004, effect-size (r) = 0.66 |
| Fig. 3E | vCA1 vs dCA1 CS+/baseline decoding accuracies during trace period (Late session) | 10 decoding iterations for each region | n-matched pseudopopulation of 537 cells from 4 vCA1 and 2 dCA1 mice | Mann-Whitney U | U = 40.5, p = 0.5, effect-size (r) = 0.16 |
| Fig. 3F | CS+/CS- decoding accuracy, vCA1 vs dCA1 | 10 decoding iterations for each region | n-matched pseudopopulation of 537 cells from 4 vCA1 and 2 dCA1 mice | Mann-Whitney U | colored-coded bar above graph shows time bins where p < 0.01 |
| Fig. 3G | vCA1 vs dCA1 CS+/CS- decoding accuracies during tone period (Early session) | 10 decoding iterations for each region | n-matched pseudopopulation of 537 cells from 4 vCA1 and 2 dCA1 mice | Mann-Whitney U | U = 52, p = 0.91, effect-size (r) = 0.03 |
| Fig. 3G | vCA1 vs dCA1 CS+/CS- decoding accuracies during tone period (Late session) | 10 decoding iterations for each region | n-matched pseudopopulation of 537 cells from 4 vCA1 and 2 dCA1 mice | Mann-Whitney U | U = 99, p < 0.001, effect-size (r) = 0.82 |
| Fig. 3G | vCA1 vs dCA1 CS+/CS- decoding accuracies during trace period (Early session) | 10 decoding iterations for each region | n-matched pseudopopulation of 537 cells from 4 vCA1 and 2 dCA1 mice | Mann-Whitney U | U = 77, p = 0.045, effect-size (r) = 0.46 |
| Fig. 3G | vCA1 vs dCA1 CS+/CS- decoding accuracies during trace period (Late session) | 10 decoding iterations for each region | n-matched pseudopopulation of 537 cells from 4 vCA1 and 2 dCA1 mice | Mann-Whitney U | U = 51, p = 0.97, effect-size (r) = 0.02 |
| Fig. 4E | Early vs Late CS+/CS- decoding accuracies during odor period (vCA1) | 10 decoding iterations for each region | n-matched pseudopopulation of 454 cells from 11 vCA1 mice | Mann-Whitney U, Bonferroni correction for n=3 | U = 0, p < 0.001, effect-size (r) = 0.85 |
| Fig. 4E | Early vs Ext CS+/CS- decoding accuracies during odor period (vCA1) | 10 decoding iterations for each region | n-matched pseudopopulation of 454 cells from 11 vCA1 mice | Mann-Whitney U, Bonferroni correction for n=3 | U = 83.5, p = 0.038, effect-size (r) = 0.57 |
| Fig. 4E | Early vs Reacq CS+/CS- decoding accuracies during odor period (vCA1) | 10 decoding iterations for each region | n-matched pseudopopulation of 454 cells from 11 vCA1 mice | Mann-Whitney U, Bonferroni correction for n=3 | U =13.5 , p = 0.019, effect-size (r) = 0.62 |
| Fig. 4E | Late vs Ext CS+/CS- decoding accuracies during odor period (vCA1) | 10 decoding iterations for each region | n-matched pseudopopulation of 454 cells from 11 vCA1 mice | Mann-Whitney U, Bonferroni correction for n=3 | U = 100, p < 0.001, effect-size (r) = 0.85 |
| Fig. 4E | Late vs Reacq CS+/CS- decoding accuracies during odor period (vCA1) | 10 decoding iterations for each region | n-matched pseudopopulation of 454 cells from 11 vCA1 mice | Mann-Whitney U, Bonferroni correction for n=3 | U = 99, p < 0.001, effect-size (r) = 0.83 |
| Fig. 4E | Ext vs Reacq CS+/CS- decoding accuracies during odor period (vCA1) | 10 decoding iterations for each region | n-matched pseudopopulation of 454 cells from 11 vCA1 mice | Mann-Whitney U, Bonferroni correction for n=3 | U = 2.5, p = 0.001, effect-size (r) = 0.80 |
| Fig. 4E | Early vs Late CS+/CS- decoding accuracies during odor period (dCA1) | 10 decoding iterations for each region | n-matched pseudopopulation of 454 cells from 4 dCA1 mice | Mann-Whitney U, Bonferroni correction for n=3 | U = 42, p = 1, effect-size (r) = 0.14 |
| Fig. 4E | Early vs Ext CS+/CS- decoding accuracies during odor period (dCA1) | 10 decoding iterations for each region | n-matched pseudopopulation of 454 cells from 4 dCA1 mice | Mann-Whitney U, Bonferroni correction for n=3 | U = 70.5, p = 0.39, effect-size (r) = 0.35 |
| Fig. 4E | Early vs Reacq CS+/CS- decoding accuracies during odor period (dCA1) | 10 decoding iterations for each region | n-matched pseudopopulation of 454 cells from 4 dCA1 mice | Mann-Whitney U, Bonferroni correction for n=3 | U = 62.5, p = 1, effect-size (r) = 0.21 |
| Fig. 4E | Late vs Ext CS+/CS- decoding accuracies during odor period (dCA1) | 10 decoding iterations for each region | n-matched pseudopopulation of 454 cells from 4 dCA1 mice | Mann-Whitney U, Bonferroni correction for n=3 | U = 79.5, p = 0.1, effect-size (r) = 0.13 |
| Fig. 4E | Late vs Reacq CS+/CS- decoding accuracies during odor period (dCA1) | 10 decoding iterations for each region | n-matched pseudopopulation of 454 cells from 4 dCA1 mice | Mann-Whitney U, Bonferroni correction for n=3 | U = 74, p = 0.22, effect-size (r) = 0.41 |
| Fig. 4E | Ext vs Reacq CS+/CS- decoding accuracies during odor period (dCA1) | 10 decoding iterations for each region | n-matched pseudopopulation of 454 cells from 4 dCA1 mice | Mann-Whitney U, Bonferroni correction for n=3 | U = 42, p = 1, effect-size (r) = 0.13 |
| Fig. 4E | Early vs Late CS+/CS- decoding accuracies during odor period (vCA1) | 10 decoding iterations for each region | n-matched pseudopopulation of 454 cells from 11 vCA1 mice | Mann-Whitney U, Bonferroni correction for n=3 | U = 0, p < 0.001, effect-size (r) = 0.85 |
| Fig. 4E | Early vs Ext CS+/CS- decoding accuracies during odor period (vCA1) | 10 decoding iterations for each region | n-matched pseudopopulation of 454 cells from 11 vCA1 mice | Mann-Whitney U, Bonferroni correction for n=3 | U = 58.5, p = 1, effect-size (r) = 0.14 |

| Figure | Variable | Unit of Comparison | n | Test | Results |
|---|---|---|---|---|---|
| Fig. 4E | Early vs Reacq CS+/CS- decoding accuracies during odor period (vCA1) | 10 decoding iterations for each region | n-matched pseudopopulation of 454 cells from 11 vCA1 mice | Mann-Whitney U, Bonferroni correction for n=3 | U = 0, p < 0.001, effect-size (r) = 0.85 |
| Fig. 4E | Late vs Ext CS+/CS- decoding accuracies during odor period (vCA1) | 10 decoding iterations for each region | n-matched pseudopopulation of 454 cells from 11 vCA1 mice | Mann-Whitney U, Bonferroni correction for n=3 | U = 0, p < 0.001, effect-size (r) = 0.85 |
| Fig. 4E | Late vs Reacq CS+/CS- decoding accuracies during odor period (vCA1) | 10 decoding iterations for each region | n-matched pseudopopulation of 454 cells from 11 vCA1 mice | Mann-Whitney U, Bonferroni correction for n=3 | U = 46.5, p = 1., effect-size (r) = 0.14 |
| Fig. 4E | Ext vs Reacq CS+/CS- decoding accuracies during odor period (vCA1) | 10 decoding iterations for each region | n-matched pseudopopulation of 454 cells from 11 vCA1 mice | Mann-Whitney U, Bonferroni correction for n=3 | U = 0, p < 0.001, effect-size (r) = 0.85 |
| Fig. 4E | Early vs Late CS+/CS- decoding accuracies during odor period (dCA1) | 10 decoding iterations for each region | n-matched pseudopopulation of 454 cells from 4 dCA1 mice | Mann-Whitney U, Bonferroni correction for n=3 | U = 0, p < 0.001, effect-size (r) = 0.85 |
| Fig. 4E | Early vs Ext CS+/CS- decoding accuracies during odor period (dCA1) | 10 decoding iterations for each region | n-matched pseudopopulation of 454 cells from 4 dCA1 mice | Mann-Whitney U, Bonferroni correction for n=3 | U = 26, p = 0.23, effect-size (r) = 0.41 |
| Fig. 4E | Early vs Reacq CS+/CS- decoding accuracies during odor period (dCA1) | 10 decoding iterations for each region | n-matched pseudopopulation of 454 cells from 4 dCA1 mice | Mann-Whitney U, Bonferroni correction for n=3 | U = 0, p < 0.001, effect-size (r) = 0.85 |
| Fig. 4E | Late vs Ext CS+/CS- decoding accuracies during odor period (dCA1) | 10 decoding iterations for each region | n-matched pseudopopulation of 454 cells from 4 dCA1 mice | Mann-Whitney U, Bonferroni correction for n=3 | U = 0, p < 0.001, effect-size (r) = 0.85 |
| Fig. 4E | Late vs Reacq CS+/CS- decoding accuracies during odor period (dCA1) | 10 decoding iterations for each region | n-matched pseudopopulation of 454 cells from 4 dCA1 mice | Mann-Whitney U, Bonferroni correction for n=3 | U = 80.5, p = 0.07, effect-size (r) = 0.52 |
| Fig. 4E | Ext vs Reacq CS+/CS- decoding accuracies during odor period (dCA1) | 10 decoding iterations for each region | n-matched pseudopopulation of 454 cells from 4 dCA1 mice | Mann-Whitney U, Bonferroni correction for n=3 | U = 0, p < 0.001, effect-size (r) = 0.85 |
| Fig. 4F | within-session Late vs within-session Reacq CS+/CS- decoding accuracies during odor period (vCA1) | 10 decoding iterations for each | n-matched pseudopopulation of 241 cells from 10 vCA1 and 4 dCA1 mice | Mann-Whitney U, Bonferroni correction for n=2 | U = 51.5, p = 1, effect-size (r) = 0.03 |
| Fig. 4F | within-session Late vs across-session Late/Reacq CS+/CS- decoding accuracies during odor period (vCA1) | 10 decoding iterations for each | n-matched pseudopopulation of 241 cells from 10 vCA1 and 4 dCA1 mice | Mann-Whitney U, Bonferroni correction for n=2 | U = 81.5, p = 0.03, effect-size (r) = 0.53 |
| Fig. 4F | within-session Reacq vs across-session Late/Reacq CS+/CS- decoding accuracies during odor period (vCA1) | 10 decoding iterations for each | n-matched pseudopopulation of 241 cells from 10 vCA1 and 4 dCA1 mice | Mann-Whitney U, Bonferroni correction for n=2 | U = 92.5, p = 0.003, effect-size (r) = 0.72 |
| Fig. 4F | within-session Late vs within-session Reacq CS+/CS- decoding accuracies during odor period (dCA1) | 10 decoding iterations for each | n-matched pseudopopulation of 241 cells from 10 vCA1 and 4 dCA1 mice | Mann-Whitney U, Bonferroni correction for n=2 | U 55= , p = 1, effect-size (r) = 0.08 |
| Fig. 4F | within-session Late vs across-session Late/Reacq CS+/CS- decoding accuracies during odor period (dCA1) | 10 decoding iterations for each | n-matched pseudopopulation of 241 cells from 10 vCA1 and 4 dCA1 mice | Mann-Whitney U, Bonferroni correction for n=2 | U = 77, p = 0.09, effect-size (r) = 0.46 |
| Fig. 4F | within-session Reacq vs across-session Late/Reacq CS+/CS- decoding accuracies during odor period (dCA1) | 10 decoding iterations for each | n-matched pseudopopulation of 241 cells from 10 vCA1 and 4 dCA1 mice | Mann-Whitney U, Bonferroni correction for n=2 | U = 96, p = 0.001, effect-size (r) = 0.78 |
| Fig. 4F | within-session Late vs within-session Reacq CS+/CS- decoding accuracies during trace period (vCA1) | 10 decoding iterations for each | n-matched pseudopopulation of 241 cells from 10 vCA1 and 4 dCA1 mice | Mann-Whitney U, Bonferroni correction for n=2 | U = 45, p = 1, effect-size (r) = 0.08 |
| Fig. 4F | within-session Late vs across-session Late/Reacq CS+/CS- decoding accuracies during trace period (vCA1) | 10 decoding iterations for each | n-matched pseudopopulation of 241 cells from 10 vCA1 and 4 dCA1 mice | Mann-Whitney U, Bonferroni correction for n=2 | U = 72, p = 0.21, effect-size (r) = 0.37 |
| Fig. 4F | within-session Reacq vs across-session Late/Reacq CS+/CS- decoding accuracies during trace period (vCA1) | 10 decoding iterations for each | n-matched pseudopopulation of 241 cells from 10 vCA1 and 4 dCA1 mice | Mann-Whitney U, Bonferroni correction for n=2 | U = 81, p = 0.042, effect-size (r) = 0.52 |
| Fig. 4F | within-session Late vs within-session Reacq CS+/CS- decoding accuracies during trace period (dCA1) | 10 decoding iterations for each | n-matched pseudopopulation of 241 cells from 10 vCA1 and 4 dCA1 mice | Mann-Whitney U, Bonferroni correction for n=2 | U = 78, p = 0.08, effect-size (r) = 0.47 |
| Fig. 4F | within-session Late vs across-session Late/Reacq CS+/CS- decoding accuracies during trace period (dCA1) | 10 decoding iterations for each | n-matched pseudopopulation of 241 cells from 10 vCA1 and 4 dCA1 mice | Mann-Whitney U, Bonferroni correction for n=2 | U = 92.5, p = 0.003, effect-size (r) = 0.72 |
| Fig. 4F | within-session Reacq vs across-session Late/Reacq CS+/CS- decoding accuracies during trace period (dCA1) | 10 decoding iterations for each | n-matched pseudopopulation of 241 cells from 10 vCA1 and 4 dCA1 mice | Mann-Whitney U, Bonferroni correction for n=2 | U = 83, p = 0.028, effect-size (r) = 0.55 |
| Fig. 4G | Across-session Early/Late vs Late/Reacq CS+/CS- decoding accuracies during dodr period (vCA1) | 10 decoding iterations for each | n-matched pseudopopulation of 241 cells from 10 vCA1 mice | Mann-Whitney U | U = 10, p = 0.003, effect-size (r) = 0.68 |
| Fig. 4G | Across-session Early/Late vs Late/Reacq CS+/CS- decoding accuracies during odor period (dCA1) | 10 decoding iterations for each | n-matched pseudopopulation of 241 cells from 4 dCA1 mice | Mann-Whitney U | U = 0, p < 0.001, effect-size (r) = 0.85 |
| Fig. 4G | Across-session Early/Late vs Late/Reacq CS+/CS- decoding accuracies during trace period (vCA1) | 10 decoding iterations for each | n-matched pseudopopulation of 241 cells from 10 vCA1 mice | Mann-Whitney U | U = 0, p < 0.001, effect-size (r) = 0.85 |
| Fig. 4G | Across-session Early/Late vs Late/Reacq CS+/CS- decoding accuracies during trace period (dCA1) | 10 decoding iterations for each | n-matched pseudopopulation of 241 cells from 4 dCA1 mice | Mann-Whitney U | U = 0, p < 0.001, effect-size (r) = 0.85 |

| Figure | Variable | Unit of Comparison | n | Test | Results |
|---|---|---|---|---|---|
| Fig. 5B | CS+/CS- decoding accuracy, vCA1 vs dCA1 | 10 decoding iterations for each | n-matched pseudopopulation of 339 cells from 7 vCA1 and 2 dCA1 mice | Mann-Whitney U | see figure for color-coded p-values |
| Fig. 5C | CS+/CS- Early decoding accuracy, vCA1 vs dCA1 | n = 19 (mean of each time bin x vs time bin y decoding result (each blue square in fig)) | n-matched pseudopopulation of 339 cells from 7 vCA1 and 2 dCA1 mice | Mann-Whitney U | U = 254, p = 0.033, effect-size (r) = 0.35 |
| Fig. 5C | CS+/CS- Late decoding accuracy, vCA1 vs dCA1 | n = 19 (mean of each time bin x vs time bin y decoding result (each blue square in fig)) | n-matched pseudopopulation of 339 cells from 7 vCA1 and 2 dCA1 mice | Mann-Whitney U | U =313 , p < 0.001., effect-size (r) = 0.63 |
| Fig. 5C | CS+/CS- Extinction decoding accuracy, vCA1 vs dCA1 | n = 19 (mean of each time bin x vs time bin y decoding result (each blue square in fig)) | n-matched pseudopopulation of 339 cells from 7 vCA1 and 2 dCA1 mice | Mann-Whitney U | U = 236.5, p = 0.11, effect-size (r) = 0.27 |
| Fig. 5C | CS+/CS- Reacquisition decoding accuracy, vCA1 vs dCA1 | n = 19 (mean of each time bin x vs time bin y decoding result (each blue square in fig)) | n-matched pseudopopulation of 339 cells from 7 vCA1 and 2 dCA1 mice | Mann-Whitney U | U = 321, p < 0.001, effect-size (r) = 0.67 |
| Fig. 6B | mean lick rate (Hz) | Trial type (CS1+ vs CS2+; Early session) | 9 mice ( 7vCA1 and 2 dCA1 mice) | Mann-Whitney U | U = 18, p = 0.94, effect-size (r) = 0 |
| Fig. 6B | mean lick rate (Hz) | Trial type (CS3- vs CS4-; Early session) | 9 mice ( 7vCA1 and 2 dCA1 mice) | Mann-Whitney U | U = 14, p = 0.58, effect-size (r) = 0.18 |
| Fig. 6B | mean lick rate (Hz) | Trial type (CS+ vs CS-; Early session) | 9 mice ( 7vCA1 and 2 dCA1 mice) | Mann-Whitney U | U = 129, p = 0.001, effect-size (r) = 0.67 |
| Fig. 6B | mean lick rate (Hz) | Trial type (CS1+ vs CS2+; Late session) | 9 mice ( 7vCA1 and 2 dCA1 mice) | Mann-Whitney U | U = 92.5, p = 0.82, effect-size (r) = 0.05 |
| Fig. 6B | mean lick rate (Hz) | Trial type (CS3- vs CS4-; Late session) | 9 mice ( 7vCA1 and 2 dCA1 mice) | Mann-Whitney U | U = 84, p = 0.53, effect-size (r) = 0.12 |
| Fig. 6B | mean lick rate (Hz) | Trial type (CS+ vs CS-; Late session) | 9 mice ( 7vCA1 and 2 dCA1 mice) | Mann-Whitney U | U = 784, p < 0.001, effect-size (r) = 0.86 |
| Fig. 6D | outcome decoding, odor period vs trace period (vCA1) | 10 decoding iterations for each | n-matched pseudopopulation of 339 cells from 7 vCA1 mice | Mann-Whitney U | U = 0, p < 0.001, effect-size (r) = 0.85 |
| Fig. 6D | outcome decoding, odor period vs trace period (dCA1) | 10 decoding iterations for each | n-matched pseudopopulation of 339 cells from 2 dCA1 mice | Mann-Whitney U | U = 0, p < 0.001, effect-size (r) = 0.85 |
| Fig. 7B | mean lick rate (Hz) | Trial type (Rew vs Shock; Early session) | 13 mice ( 10 vCA1 and 3 dCA1 mice) | Mann-Whitney U, Bonferroni correction for n=2 | U = 133, p = 0.028, effect-size (r) = 0.49 |
| Fig. 7B | mean lick rate (Hz) | Trial type (Rew vs CS-; Early session) | 13 mice ( 10 vCA1 and 3 dCA1 mice) | Mann-Whitney U, Bonferroni correction for n=2 | U = 13, p < 0.001, effect-size (r) = 0.71 |
| Fig. 7B | mean lick rate (Hz) | Trial type (Shock vs CS-; Early session) | 13 mice ( 10 vCA1 and 3 dCA1 mice) | Mann-Whitney U, Bonferroni correction for n=2 | U = 12.5, p <0.001, effect-size (r) = 0.72 |
| Fig. 7B | mean lick rate (Hz) | Trial type (Rew vs Shock; Late session) | 13 mice ( 10 vCA1 and 3 dCA1 mice) | Mann-Whitney U, Bonferroni correction for n=2 | U = 256, p = < 0.001, effect-size (r) = 1.7 |
| Fig. 7B | mean lick rate (Hz) | Trial type (Rew vs CS-; Late session) | 13 mice ( 10 vCA1 and 3 dCA1 mice) | Mann-Whitney U, Bonferroni correction for n=2 | U = 0, p < 0.001, effect-size (r) = 0.85 |
| Fig. 7B | mean lick rate (Hz) | Trial type (Shock vs CS-; Late session) | 13 mice ( 10 vCA1 and 3 dCA1 mice) | Mann-Whitney U, Bonferroni correction for n=2 | U = 0, p = 1, effect-size (r) = 0.56 |
| Fig. 7E | CS+Rew Decoding Accuracy, Early vs Late, odor period (vCA1) | 10 decoding iterations for each | n-matched pseudopopulation of 444 cells from 10 vCA1 mice | Fisher's Exact | p < 0.001, effect-size (odds ratio) = 0.23 |
| Fig. 7E | CS+Shock Decoding Accuracy, Early vs Late, odor period (vCA1) | 10 decoding iterations for each | n-matched pseudopopulation of 444 cells from 10 vCA1 mice | Fisher's Exact | p < 0.001, effect-size (odds ratio) = 0.24 |
| Fig. 7E | CS- Decoding Accuracy, Early vs Late, odor period (vCA1) | 10 decoding iterations for each | n-matched pseudopopulation of 444 cells from 10 vCA1 mice | Fisher's Exact | p = 0.14, effect-size (odds ratio) = 0.72 |
| Fig. 7E | CS+Rew Decoding Accuracy, Early vs Late,trace period (vCA1) | 10 decoding iterations for each | n-matched pseudopopulation of 444 cells from 10 vCA1 mice | Fisher's Exact | p < 0.001, effect-size (odds ratio) = 0.26 |
| Fig. 7E | CS+Shock Decoding Accuracy, Early vs Late, trace period (vCA1) | 10 decoding iterations for each | n-matched pseudopopulation of 444 cells from 10 vCA1 mice | Fisher's Exact | p < 0.001, effect-size (odds ratio) = 0.40 |
| Fig. 7E | CS- Decoding Accuracy, Early vs Late, trace period (vCA1) | 10 decoding iterations for each | n-matched pseudopopulation of 444 cells from 10 vCA1 mice | Fisher's Exact | p = 0.24, effect-size (odds ratio) = 0.76 |
| Fig. 7F | CS+Rew Decoding Accuracy, Early vs Late,odor period (dCA1) | 10 decoding iterations for each | n-matched pseudopopulation of 444 cells from 3 dCA1 mice | Fisher's Exact | p = 0.003, effect-size (odds ratio) = 0.18 |
| Fig. 7F | CS+Shock Decoding Accuracy, Early vs Late, odor period (dCA1) | 10 decoding iterations for each | n-matched pseudopopulation of 444 cells from 3 dCA1 mice | Fisher's Exact | p = 0.32, effect-size (odds ratio) = 0.63 |
| Fig. 7F | CS- Decoding Accuracy, Early vs Late, odor period (dCA1) | 10 decoding iterations for each | n-matched pseudopopulation of 444 cells from 3 dCA1 mice | Fisher's Exact | p = 0.73, effect-size (odds ratio) = 0.88 |
| Fig. 7F | CS+Rew Decoding Accuracy, Early vs Late,trace period (dCA1) | 10 decoding iterations for each | n-matched pseudopopulation of 444 cells from 3 dCA1 mice | Fisher's Exact | p < 0.001, effect-size (odds ratio) = 0.17 |
| Fig. 7F | CS+Shock Decoding Accuracy, Early vs Late, trace period (dCA1) | 10 decoding iterations for each | n-matched pseudopopulation of 444 cells from 3 dCA1 mice | Fisher's Exact | p = 0.003, effect-size (odds ratio) = 0.48 |
| Fig. 7F | CS- Decoding Accuracy, Early vs Late, trace period (dCA1) | 10 decoding iterations for each | n-matched pseudopopulation of 444 cells from 3 dCA1 mice | Fisher's Exact | p = 0.003, effect-size (odds ratio) = 0.52 |
| Fig. 7H | CS+Rew/CS- Late decoding accuracy, vCA1 vs dCA1 | n = 19 (mean of each time bin x vs time bin y decoding result (each blue square in fig)) | n-matched pseudopopulation of 444 cells from 10 vCA1 and 3 dCA1 mice | Mann-Whitney U | U = 346, p <0.001, effect-size (r) = 0.79 |

| Figure | Variable | Unit of Comparison | n | Test | Results |
|---|---|---|---|---|---|
| Fig. 7J | CS+Shock/CS- Late decoding accuracy, vCA1 vs dCA1 | n = 19 (mean of each time bin x vs time bin y decoding result (each blue square in fig)) | n-matched pseudopopulation of 444 cells from 10 vCA1 and 3 dCA1 mice | Mann-Whitney U | U = 326, p < 0.001, effect-size (r) = 0.69 |
| Fig. 7L | mean lick rate (Hz) | Trial type (Rew vs Shock; Early Reversal session) | 13 mice ( 10 vCA1 and 3 dCA1 mice) | Mann-Whitney U, Bonferroni correction for n=2 | U = 82, p = 1, effect-size (r) = 0.02 |
| Fig. 7L | mean lick rate (Hz) | Trial type (Rew vs CS-; Early Reversal session) | 13 mice ( 10 vCA1 and 3 dCA1 mice) | Mann-Whitney U, Bonferroni correction for n=2 | U = 21.5, p = 0.003, effect-size (r) = 0.62 |
| Fig. 7L | mean lick rate (Hz) | Trial type (Shock vs CS-; Early Reversal session) | 13 mice ( 10 vCA1 and 3 dCA1 mice) | Mann-Whitney U, Bonferroni correction for n=2 | U = 10, p <0.001, effect-size (r) = 0.74 |
| Fig. 7L | mean lick rate (Hz) | Trial type (Rew vs Shock; Late Reversal session) | 13 mice ( 10 vCA1 and 3 dCA1 mice) | Mann-Whitney U, Bonferroni correction for n=2 | U = 255, p = < 0.001, effect-size (r) = 1.7 |
| Fig. 7L | mean lick rate (Hz) | Trial type (Rew vs CS-; Late Reversal session) | 13 mice ( 10 vCA1 and 3 dCA1 mice) | Mann-Whitney U, Bonferroni correction for n=2 | U = 3, p < 0.001, effect-size (r) = 0.81 |
| Fig. 7L | mean lick rate (Hz) | Trial type (Shock vs CS-; Late Reversal session) | 13 mice ( 10 vCA1 and 3 dCA1 mice) | Mann-Whitney U, Bonferroni correction for n=2 | U = 110, p = 1, effect-size (r) = 0.26 |
| Fig. 7M | Odor A/baseline vs odor B/baseline decoding accuracies across reversal training, odor period (vCA1) | 10 decoding iterations for each | n-matched pseudopopulation of 281 cells from 10 vCA1 mice | Mann-Whitney U, Bonferroni correction for n=2 | U = 41.5, p = 1, effect-size (r) = 0.14 |
| Fig. 7M | Odor A/baseline vs odor C/baseline decoding accuracies across reversal training, odor period (vCA1) | 10 decoding iterations for each | n-matched pseudopopulation of 281 cells from 10 vCA1 mice | Mann-Whitney U, Bonferroni correction for n=2 | U = 0, p < 0.001, effect-size (r) = 0.85 |
| Fig. 7M | Odor B/baseline vs odor C/baseline decoding accuracies across reversal training, odor period (vCA1) | 10 decoding iterations for each | n-matched pseudopopulation of 281 cells from 10 vCA1 mice | Mann-Whitney U, Bonferroni correction for n=2 | U = 0, p < 0.001, effect-size (r) = 0.85 |
| Fig. 7M | Odor A/baseline vs odor B/baseline decoding accuracies across reversal training, odor period (dCA1) | 10 decoding iterations for each | n-matched pseudopopulation of 281 cells from 3 dCA1 mice | Mann-Whitney U, Bonferroni correction for n=2 | U = 23.5, p = 0.1, effect-size (r) = 0.45 |
| Fig. 7M | Odor A/baseline vs odor C/baseline decoding accuracies across reversal training, odor period (dCA1) | 10 decoding iterations for each | n-matched pseudopopulation of 281 cells from 3 dCA1 mice | Mann-Whitney U, Bonferroni correction for n=2 | U = 58, p = 1, effect-size (r) = 0.14 |
| Fig. 7M | Odor B/baseline vs odor C/baseline decoding accuracies across reversal training, odor period (dCA1) | 10 decoding iterations for each | n-matched pseudopopulation of 281 cells from 3 dCA1 mice | Mann-Whitney U, Bonferroni correction for n=2 | U =28.5 , p = 0.22, effect-size (r) = 0.36 |
| Fig. 7N | CS+Rew/baseline vs CS+Shock/baseline decoding accuracies across reversal training, trace period (vCA1) | 10 decoding iterations for each | n-matched pseudopopulation of 281 cells from 10 vCA1 mice | Mann-Whitney U, Bonferroni correction for n=2 | U = 100, p < 0.001, effect-size (r) = 0.85 |
| Fig. 7N | CS+Rew/baseline vs CS-/baseline decoding accuracies across reversal training, trace period (vCA1) | 10 decoding iterations for each | n-matched pseudopopulation of 281 cells from 10 vCA1 mice | Mann-Whitney U, Bonferroni correction for n=2 | U = 0, p < 0.001, effect-size (r) = 0.85 |
| Fig. 7N | CS+Shock/baseline vs CS-/baseline decoding accuracies across reversal training, trace period (vCA1) | 10 decoding iterations for each | n-matched pseudopopulation of 281 cells from 10 vCA1 mice | Mann-Whitney U, Bonferroni correction for n=2 | U = 55, p = 1, effect-size (r) = 0.08 |
| Fig. 7N | CS+Rew/baseline vs CS+Shock/baseline decoding accuracies across reversal training, trace period (dCA1) | 10 decoding iterations for each | n-matched pseudopopulation of 281 cells from 3 dCA1 mice | Mann-Whitney U, Bonferroni correction for n=2 | U = 100, p < 0.001, effect-size (r) = 0.85 |
| Fig. 7N | CS+Rew/baseline vs CS-/baseline decoding accuracies across reversal training, trace period (dCA1) | 10 decoding iterations for each | n-matched pseudopopulation of 281 cells from 3 dCA1 mice | Mann-Whitney U, Bonferroni correction for n=2 | U = 0, p < 0.001, effect-size (r) = 0.85 |
| Fig. 7N | CS+Shock/baseline vs CS-/baseline decoding accuracies across reversal training, trace period (dCA1) | 10 decoding iterations for each | n-matched pseudopopulation of 281 cells from 3 dCA1 mice | Mann-Whitney U, Bonferroni correction for n=2 | U = 49, p = 1, effect-size (r) = 0.17 |
| Fig. 7O | CS+Rew/baseline vs CS+Shock/baseline decoding accuracies, Late Reversal, odor period (vCA1) | 10 decoding iterations for each | n-matched pseudopopulation of 444 cells from 10 vCA1 mice | Mann-Whitney U, Bonferroni correction for n=2 | U = 100, p < 0.001, effect-size (r) = 0.85 |
| Fig. 7O | CS+Rew/baseline vs CS-/baseline decoding accuracies, Late Reversal, odor period (vCA1) | 10 decoding iterations for each | n-matched pseudopopulation of 444 cells from 10 vCA1 mice | Mann-Whitney U, Bonferroni correction for n=2 | U = 100, p < 0.001, effect-size (r) = 0.85 |
| Fig. 7O | CS+Shock/baseline vs CS-/baseline decoding accuracies, Late Reversal, odor period (vCA1) | 10 decoding iterations for each | n-matched pseudopopulation of 444 cells from 10 vCA1 mice | Mann-Whitney U, Bonferroni correction for n=2 | U = 97, p < 0.001, effect-size (r) = 0.79 |
| Fig. 7O | CS+Rew/baseline vs CS+Shock/baseline decoding accuracies, Late Reversal, odor period (dCA1) | 10 decoding iterations for each | n-matched pseudopopulation of 444 cells from 3 dCA1 mice | Mann-Whitney U, Bonferroni correction for n=2 | U = 17.5, p = 0.03, effect-size (r) = 0.55 |
| Fig. 7O | CS+Rew/baseline vs CS-/baseline decoding accuracies, Late Reversal, odor period (dCA1) | 10 decoding iterations for each | n-matched pseudopopulation of 444 cells from 3 dCA1 mice | Mann-Whitney U, Bonferroni correction for n=2 | U = 27.5, p = 0.19, effect-size (r) = 0.38 |
| Fig. 7O | CS+Shock/baseline vs CS-/baseline decoding accuracies, Late Reversal, odor period (dCA1) | 10 decoding iterations for each | n-matched pseudopopulation of 444 cells from 3 dCA1 mice | Mann-Whitney U, Bonferroni correction for n=2 | U = 55.5, p = 1, effect-size (r) = 0.09 |

| Figure | Variable | Unit of Comparison | n | Test | Results |
|---|---|---|---|---|---|
| Fig. 7O | CS+Rew/baseline vs CS+Shock/baseline decoding accuracies, Late Reversal, trace period (vCA1) | 10 decoding iterations for each | n-matched pseudopopulation of 444 cells from 10 vCA1 mice | Mann-Whitney U, Bonferroni correction for n=2 | U = 100, p < 0.001, effect-size (r) = 0.85 |
| Fig. 7O | CS+Rew/baseline vs CS-/baseline decoding accuracies, Late Reversal, trace period (vCA1) | 10 decoding iterations for each | n-matched pseudopopulation of 444 cells from 10 vCA1 mice | Mann-Whitney U, Bonferroni correction for n=2 | U = 100, p < 0.001, effect-size (r) = 0.85 |
| Fig. 7O | CS+Shock/baseline vs CS-/baseline decoding accuracies, Late Reversal, trace period (vCA1) | 10 decoding iterations for each | n-matched pseudopopulation of 444 cells from 10 vCA1 mice | Mann-Whitney U, Bonferroni correction for n=2 | U = 22.5, p = 0.08, effect-size (r) = 0.46 |
| Fig. 7O | CS+Rew/baseline vs CS+Shock/baseline decoding accuracies, Late Reversal, trace period (dCA1) | 10 decoding iterations for each | n-matched pseudopopulation of 444 cells from 3 dCA1 mice | Mann-Whitney U, Bonferroni correction for n=2 | U = 100, p < 0.001, effect-size (r) = 0.85 |
| Fig. 7O | CS+Rew/baseline vs CS-/baseline decoding accuracies, Late Reversal, trace period (dCA1) | 10 decoding iterations for each | n-matched pseudopopulation of 444 cells from 3 dCA1 mice | Mann-Whitney U, Bonferroni correction for n=2 | U = 100, p < 0.001, effect-size (r) = 0.85 |
| Fig. 7O | CS+Shock/baseline vs CS-/baseline decoding accuracies, Late Reversal, trace period (dCA1) | 10 decoding iterations for each | n-matched pseudopopulation of 444 cells from 3 dCA1 mice | Mann-Whitney U, Bonferroni correction for n=2 | U = 23.5, p = 0.095, effect-size (r) = 0.45 |
| Fig. S2C | CS+ responsive cells during odor period, Early vs Late (vCA1) | total combined cells from 11 vCA1 mice | see figure for exact cell numbers | Fisher's Exact | p = 0.003, effect-size (odds ratio) = 2.02 |
| Fig. S2C | CS- responsive cells during odor period, Early vs Late (vCA1) | total combined cells from 11 vCA1 mice | see figure for exact cell numbers | Fisher's Exact | p = 0.13, effect-size (odds ratio) = 0.6 |
| Fig. S2C | CS+ responsive cells during trace period, Early vs Late (vCA1) | total combined cells from 11 vCA1 mice | see figure for exact cell numbers | Fisher's Exact | p < 0.001, effect-size (odds ratio) = 4.56 |
| Fig. S2C | CS- responsive cells during trace period, Early vs Late (vCA1) | total combined cells from 11 vCA1 mice | see figure for exact cell numbers | Fisher's Exact | p = 0.055, effect-size (odds ratio) = 6.96 |
| Fig. S2C | CS+ responsive cells during odor period, Early vs Late (dCA1) | total combined cells from 4 dCA1 mice | see figure for exact cell numbers | Fisher's Exact | p < 0.001, effect-size (odds ratio) = 2.7 |
| Fig. S2C | CS- responsive cells during odor period, Early vs Late (dCA1) | total combined cells from 4 dCA1 mice | see figure for exact cell numbers | Fisher's Exact | p = 0.38, effect-size (odds ratio) = 0.79 |
| Fig. S2C | CS+ responsive cells during trace period, Early vs Late (dCA1) | total combined cells from 4 dCA1 mice | see figure for exact cell numbers | Fisher's Exact | p < 0.001, effect-size (odds ratio) = 8.24 |
| Fig. S2C | CS- responsive cells during trace period, Early vs Late (dCA1) | total combined cells from 4 dCA1 mice | see figure for exact cell numbers | Fisher's Exact | p = 0.004, effect-size (odds ratio) = 11.3 |
| Fig. S2D | CS+/baseline (upper) or CS-/baseline (lower) decoding accuracies, Pre vs Late sessions | 10 decoding iterations for each | n-matched pseudopopulation of 454 cells from 11 vCA1 or 5 dCA1 mice | Mann-Whitney U | color-coded bars above graph show time bins where p < 0.01 |
| Fig. S2G | CS+/CS- decoding accuracy, odor period | pre vs post 'aha' point | 11 vCA1 mice | Mann-Whitney U | U = 10 , p = .003, effect-size (r) = 0.68 |
| Fig. S2G | CS+/CS- decoding accuracy, trace period | pre vs post 'aha' point | 11 vCA1 mice | Mann-Whitney U | U = 0 , p < .001, effect-size (r) = 0.84 |
| Fig. S2G | CS+/CS- decoding accuracy, odor period | pre vs post 'aha' point | 4 dCA1 mice | Mann-Whitney U | U = 27, p = 0.089., effect-size (r) = 0.39 |
| Fig. S2G | CS+/CS- decoding accuracy, trace period | pre vs post 'aha' point | 4 dCA1 mice | Mann-Whitney U | U = 19 , p = .021., effect-size (r) = 0.52. |
| Fig. S2H | Early vs Late linear regression of decoding accuracy and lick rate | Early vs Late sessions | 10 vCA1 mice | T-test | U = 1.86, p = 0.079, effect-size (Cohen's d) = 0.88 |
| Fig. S2H | Early vs Late linear regression of decoding accuracy and lick rate | Early vs Late sessions | 5 dCA1 mice | T-test | U = 1.51 p = 0.17, effect-size (Cohen's d) = 1.07 |
| Fig. S2I | odor period vs trace period decoding accuracy vs chance | 10 decoding iterations | pseudopopulation of 454 cells from 11 vCA1 mice | Wilcoxon | W = 0, p = 0.005, effect-size (r) = 0.85 |
| Fig. S2I | odor period vs trace period decoding accuracy vs chance | 10 decoding iterations | pseudopopulation of 454 cells from 5 dCA1 mice | Wilcoxon | W = 0, p = 0.005, effect-size (r) = 0.85 |
| Fig. S3C | CS+/baseline decoding accuracies during tone period, Early vs Late session (vCA1) | 10 decoding iterations for each session | n-matched pseudopopulation of 537 cells from 4 vCA1 mice | Mann-Whitney U | U = 24, p = 0.054, effect-size (r) = 0.44 |
| Fig. S3C | CS+/baseline decoding accuracies during tone period, Early vs Late session (dCA1) | 10 decoding iterations for each session | n-matched pseudopopulation of 537 cells from 2 dCA1 mice | Mann-Whitney U | U = 40, p = 0.47, effect-size (r) = 0.17 |
| Fig. S3D | CS-/baseline decoding accuracies during tone period, Early vs Late session (vCA1) | 10 decoding iterations for each session | n-matched pseudopopulation of 537 cells from 4 vCA1 mice | Mann-Whitney U | U = 47, p = 0.85, effect-size (r) = 0.05 |
| Fig. S3D | CS-/baseline decoding accuracies during tone period, Early vs Late session (dCA1) | 10 decoding iterations for each session | n-matched pseudopopulation of 537 cells from 2 dCA1 mice | Mann-Whitney U | U = 29.5, p = 0.13, effect-size (r) = 0.35 |
| Fig. S3E | mean lick rate (Hz) | Trial type (Early session) | 3 vCA1 mice (non-learners only) | Mann-Whitney U | U = 4, p = 1, effect-size (r) = 0.0 |
| Fig. S3E | mean lick rate (Hz) | Trial type (Late session) | 3 vCA1 mice (non-learners only) | Mann-Whitney U | U = 2, p = 0.38, effect-size (r) = 0.37 |
| Fig. S3F | CS+/CS- decoding accuracy. Early vs Late | 10 decoding iterations for each session | n-matched pseudopopulation of 71 cells from 4 vCA1 (learner), 3 vCA1 (nonlearner), or 2 dCA1 (learner) mice | Mann-Whitney U | color-coded bars above graph show time bins where p < 0.01 |

| Figure | Variable | Unit of Comparison | n | Test | Results |
|---|---|---|---|---|---|
| Fig. S3G | CS+/CS- decoding accuracy, tone period, vCA1 nonlearners vs vCA1 learners (Early) | 10 decoding iterations for each | n-matched pseudopopulation of 71 cells from 4 vCA1 (learner), 3 vCA1 (nonlearner), or 2 dCA1 (learner) mice | Mann-Whitney U, Bonferroni correction for n=2 | U = 30, p = 0.28, effect-size (r) = 0.34 |
| Fig. S3G | CS+/CS- decoding accuracy, tone period, vCA1 nonlearners vs dCA1 learners (Early) | 10 decoding iterations for each | n-matched pseudopopulation of 71 cells from 4 vCA1 (learner), 3 vCA1 (nonlearner), or 2 dCA1 (learner) mice | Mann-Whitney U, Bonferroni correction for n=2 | U = 70, p = 0.27, effect-size (r) = 0.34 |
| Fig. S3G | CS+/CS- decoding accuracy, tone period, vCA1 learners vs dCA1 learners (Early) | 10 decoding iterations for each | n-matched pseudopopulation of 71 cells from 4 vCA1 (learner), 3 vCA1 (nonlearner), or 2 dCA1 (learner) mice | Mann-Whitney U, Bonferroni correction for n=2 | U = 83, p = 0.028, effect-size (r) = 0.56 |
| Fig. S3G | CS+/CS- decoding accuracy, tone period, vCA1 nonlearners vs vCA1 learners (Late) | 10 decoding iterations for each | n-matched pseudopopulation of 71 cells from 4 vCA1 (learner), 3 vCA1 (nonlearner), or 2 dCA1 (learner) mice | Mann-Whitney U, Bonferroni correction for n=2 | U = 3, p < 0.001, effect-size (r) = 0.79 |
| Fig. S3G | CS+/CS- decoding accuracy, tone period, vCA1 nonlearners vs dCA1 learners (Late) | 10 decoding iterations for each | n-matched pseudopopulation of 71 cells from 4 vCA1 (learner), 3 vCA1 (nonlearner), or 2 dCA1 (learner) mice | Mann-Whitney U, Bonferroni correction for n=2 | U = 29.5, p = 0.26, effect-size (r) = 0.35 |
| Fig. S3G | CS+/CS- decoding accuracy, tone period, vCA1 learners vs dCA1 learners (Late) | 10 decoding iterations for each | n-matched pseudopopulation of 71 cells from 4 vCA1 (learner), 3 vCA1 (nonlearner), or 2 dCA1 (learner) mice | Mann-Whitney U, Bonferroni correction for n=2 | U = 92, p = 0.003, effect-size (r) = 0.71 |
| Fig. S3G | CS+/CS- decoding accuracy, trace period, vCA1 nonlearners vs vCA1 learners (Early) | 10 decoding iterations for each | n-matched pseudopopulation of 71 cells from 4 vCA1 (learner), 3 vCA1 (nonlearner), or 2 dCA1 (learner) mice | Mann-Whitney U, Bonferroni correction for n=2 | U = 58.5, p = 0.1, effect-size (r) = 0.14 |
| Fig. S3G | CS+/CS- decoding accuracy, trace period, vCA1 nonlearners vs dCA1 learners (Early) | 10 decoding iterations for each | n-matched pseudopopulation of 71 cells from 4 vCA1 (learner), 3 vCA1 (nonlearner), or 2 dCA1 (learner) mice | Mann-Whitney U, Bonferroni correction for n=2 | U = 36.5, p = 0.65, effect-size (r) = 0.23 |
| Fig. S3G | CS+/CS- decoding accuracy, trace period, vCA1 learners vs dCA1 learners (Early) | 10 decoding iterations for each | n-matched pseudopopulation of 71 cells from 4 vCA1 (learner), 3 vCA1 (nonlearner), or 2 dCA1 (learner) mice | Mann-Whitney U, Bonferroni correction for n=2 | U = 30.5, p = 0.3, effect-size (r) = 0.33 |
| Fig. S3G | CS+/CS- decoding accuracy, trace period, vCA1 nonlearners vs vCA1 learners (Late) | 10 decoding iterations for each | n-matched pseudopopulation of 71 cells from 4 vCA1 (learner), 3 vCA1 (nonlearner), or 2 dCA1 (learner) mice | Mann-Whitney U, Bonferroni correction for n=2 | U = 1, p < 0.001, effect-size (r) = 0.83 |
| Fig. S3G | CS+/CS- decoding accuracy, trace period, vCA1 nonlearners vs dCA1 learners (Late) | 10 decoding iterations for each | n-matched pseudopopulation of 71 cells from 4 vCA1 (learner), 3 vCA1 (nonlearner), or 2 dCA1 (learner) mice | Mann-Whitney U, Bonferroni correction for n=2 | U = 11, p = 0.007, effect-size (r) = 0.66 |
| Fig. S3G | CS+/CS- decoding accuracy, trace period, vCA1 learners vs dCA1 learners (Late) | 10 decoding iterations for each | n-matched pseudopopulation of 71 cells from 4 vCA1 (learner), 3 vCA1 (nonlearner), or 2 dCA1 (learner) mice | Mann-Whitney U, Bonferroni correction for n=2 | U = 83, p = 0.028, effect-size (r) = 0.56 |
| Fig. S4A | CS+/baseline vs CS-/baseline decoding accuracies | 10 decoding iterations for each trial type | n-matched pseudopopulation of 454 cells from 11 vCA1 or 4 dCA1 mice | Mann-Whitney U | color-coded bars above graph show time bins where p < 0.01 |
| Fig. S4B | CS+/baseline decoding accuracies, odor period, Early vs Late, vCA1 | 10 decoding iterations for each session | n-matched pseudopopulation of 454 cells from 11 vCA1 mice | Mann-Whitney U, bonferroni correction for n=3 | U = 0, p < 0.001, effect-size (r) = 0.85 |
| Fig. S4B | CS+/baseline decoding accuracies, odor period, Early vs Ext2, vCA1 | 10 decoding iterations for each session | n-matched pseudopopulation of 454 cells from 11 vCA1 mice | Mann-Whitney U, bonferroni correction for n=3 | U = 27.5, p = 0.29, effect-size (r) = 0.38 |
| Fig. S4B | CS+/baseline decoding accuracies, odor period, Early vs Reacquisition, vCA1 | 10 decoding iterations for each session | n-matched pseudopopulation of 454 cells from 11 vCA1 mice | Mann-Whitney U, bonferroni correction for n=3 | U = 3, p = 0.001, effect-size (r) = 0.79 |
| Fig. S4B | CS+/baseline decoding accuracies, odor period, Late vs Ext2, vCA1 | 10 decoding iterations for each session | n-matched pseudopopulation of 454 cells from 11 vCA1 mice | Mann-Whitney U, bonferroni correction for n=3 | U = 100, p < 0.001, effect-size (r) = 0.85 |
| Fig. S4B | CS+/baseline decoding accuracies, odor period, Late vs Reacquisition, vCA1 | 10 decoding iterations for each session | n-matched pseudopopulation of 454 cells from 11 vCA1 mice | Mann-Whitney U, bonferroni correction for n=3 | U = 79.5, p = 0.085, effect-size (r) = 0.5 |
| Fig. S4B | CS+/baseline decoding accuracies, odor period, Ext2 vs Reacquisition, vCA1 | 10 decoding iterations for each session | n-matched pseudopopulation of 454 cells from 11 vCA1 mice | Mann-Whitney U, bonferroni correction for n=3 | U = 8, p = 0.005, effect-size (r) = 0.71 |
| Fig. S4B | CS+/baseline decoding accuracies, odor period, Early vs Late, dCA1 | 10 decoding iterations for each session | n-matched pseudopopulation of 454 cells from 4 dCA1 mice | Mann-Whitney U, bonferroni correction for n=3 | U = 42.5, p = 1, effect-size (r) = 0.13 |
| Fig. S4B | CS+/baseline decoding accuracies, odor period, Early vs Ext2, dCA1 | 10 decoding iterations for each session | n-matched pseudopopulation of 454 cells from 4 dCA1 mice | Mann-Whitney U, bonferroni correction for n=3 | U = 44.5, p = 1, effect-size (r) = 0.09 |
| Fig. S4B | CS+/baseline decoding accuracies, odor period, Early vs Reacquisition, dCA1 | 10 decoding iterations for each session | n-matched pseudopopulation of 454 cells from 4 dCA1 mice | Mann-Whitney U, bonferroni correction for n=3 | U = 33, p = 0.64, effect-size (r) = 0.29 |
| Fig. S4B | CS+/baseline decoding accuracies, odor period, Late vs Ext2, dCA1 | 10 decoding iterations for each session | n-matched pseudopopulation of 454 cells from 4 dCA1 mice | Mann-Whitney U, bonferroni correction for n=3 | U = 57, p = 1, effect-size (r) = 0.12 |
| Fig. S4B | CS+/baseline decoding accuracies, odor period, Late vs Reacquisition, dCA1 | 10 decoding iterations for each session | n-matched pseudopopulation of 454 cells from 4 dCA1 mice | Mann-Whitney U, bonferroni correction for n=3 | U = 40.5, p = 1, effect-size (r) = 0.16 |
| Fig. S4B | CS+/baseline decoding accuracies, odor period, Ext2 vs Reacquisition, dCA1 | 10 decoding iterations for each session | n-matched pseudopopulation of 454 cells from 4 dCA1 mice | Mann-Whitney U, bonferroni correction for n=3 | U = 31, p = 0.29, effect-size (r) = 0.38 |
| Fig. S4C | CS+/baseline decoding accuracies, trace period, Early vs Late, vCA1 | 10 decoding iterations for each session | n-matched pseudopopulation of 454 cells from 11 vCA1 mice | Mann-Whitney U, bonferroni correction for n=3 | U = 0, p < 0.001, effect-size (r) = 0.85 |

| Figure | Variable | Unit of Comparison | n | Test | Results |
|---|---|---|---|---|---|
| Fig. S4C | CS+/baseline decoding accuracies, trace period, Early vs Ext2, vCA1 | 10 decoding iterations for each session | n-matched pseudopopulation of 454 cells from 11 vCA1 mice | Mann-Whitney U, bonferroni correction for n=3 | U = 96, p = 0.002, effect-size (r) = 0.78 |
| Fig. S4C | CS+/baseline decoding accuracies, trace period, Early vs Reacquisition, vCA1 | 10 decoding iterations for each session | n-matched pseudopopulation of 454 cells from 11 vCA1 mice | Mann-Whitney U, bonferroni correction for n=3 | U = 0, p < 0.001, effect-size (r) = 0.85 |
| Fig. S4C | CS+/baseline decoding accuracies, trace period, Late vs Ext2, vCA1 | 10 decoding iterations for each session | n-matched pseudopopulation of 454 cells from 11 vCA1 mice | Mann-Whitney U, bonferroni correction for n=3 | U = 100, p < 0.001, effect-size (r) = 0.85 |
| Fig. S4C | CS+/baseline decoding accuracies, trace period, Late vs Reacquisition, vCA1 | 10 decoding iterations for each session | n-matched pseudopopulation of 454 cells from 11 vCA1 mice | Mann-Whitney U, bonferroni correction for n=3 | U = 46.5, p = 1, effect-size (r) = 0.06 |
| Fig. S4C | CS+/baseline decoding accuracies, trace period, Ext2 vs Reacquisition, vCA1 | 10 decoding iterations for each session | n-matched pseudopopulation of 454 cells from 11 vCA1 mice | Mann-Whitney U, bonferroni correction for n=3 | U = 0, p < 0.001, effect-size (r) = 0.85 |
| Fig. S4C | CS+/baseline decoding accuracies, trace period, Early vs Late, dCA1 | 10 decoding iterations for each session | n-matched pseudopopulation of 454 cells from 4 dCA1 mice | Mann-Whitney U, bonferroni correction for n=3 | U = 0, p < 0.001, effect-size (r) = 0.85 |
| Fig. S4C | CS+/baseline decoding accuracies, trace period, Early vs Ext2, dCA1 | 10 decoding iterations for each session | n-matched pseudopopulation of 454 cells from 4 dCA1 mice | Mann-Whitney U, bonferroni correction for n=3 | U = 39.5, p = 1, effect-size (r) = 0.18 |
| Fig. S4C | CS+/baseline decoding accuracies, trace period, Early vs Reacquisition, dCA1 | 10 decoding iterations for each session | n-matched pseudopopulation of 454 cells from 4 dCA1 mice | Mann-Whitney U, bonferroni correction for n=3 | U = 0, p < 0.001, effect-size (r) = 0.85 |
| Fig. S4C | CS+/baseline decoding accuracies, trace period, Late vs Ext2, dCA1 | 10 decoding iterations for each session | n-matched pseudopopulation of 454 cells from 4 dCA1 mice | Mann-Whitney U, bonferroni correction for n=3 | U = 100, p < 0.001, effect-size (r) = 0.85 |
| Fig. S4C | CS+/baseline decoding accuracies, trace period, Late vs Reacquisition, dCA1 | 10 decoding iterations for each session | n-matched pseudopopulation of 454 cells from 4 dCA1 mice | Mann-Whitney U, bonferroni correction for n=3 | U = 66, p = 0.72, effect-size (r) = 0.27 |
| Fig. S4C | CS+/baseline decoding accuracies, trace period, Ext2 vs Reacquisition, dCA1 | 10 decoding iterations for each session | n-matched pseudopopulation of 454 cells from 4 dCA1 mice | Mann-Whitney U, bonferroni correction for n=3 | U = 0, p < 0.001, effect-size (r) = 0.85 |
| Fig. S5A | vCA1 vs dCA1, Late session, CS+ trials | Mean population activity during even trials (normalized to odd trial peak amplitude), 2-4 sec before/after time of odd trial peak | 16 time bins | Mann-Whitney U | U = 210, p = 0.002, effect-size (r) = 0.55 |
| Fig. S5B | vCA1 vs dCA1, Reacquisition session, CS+ trials | Mean population activity during even trials (normalized to odd trial peak amplitude), 2-4 sec before/after time of odd trial peak | 16 time bins | Mann-Whitney U | U = 199, p = 0.008, effect-size (r) = 0.47 |
| Fig. S6B | CS+/CS- vs CS1+/CS2+, baseline period, Late session (vCA1) | Euclidean distance between MDS values | 10 MDS runs | Mann-Whitney U, bonferroni correction for n=2 | U = 37, p = 0.69, effect-size (r) = 0.22 |
| Fig. S6B | CS+/CS- vs CS3-/CS4-, baseline period, Late session (vCA1) | Euclidean distance between MDS values | 10 MDS runs | Mann-Whitney U, bonferroni correction for n=2 | U = 52, p = 1, effect-size (r) = 0.03 |
| Fig. S6B | CS1+/CS2+ vs CS3-/CS4-, baseline period, Late session (vCA1) | Euclidean distance between MDS values | 10 MDS runs | Mann-Whitney U, bonferroni correction for n=2 | U = 62, p = 0.77, effect-size (r) = 0.2 |
| Fig. S6B | CS+/CS- vs CS1+/CS2+, odor period, Late session (vCA1) | Euclidean distance between MDS values | 10 MDS runs | Mann-Whitney U, bonferroni correction for n=2 | U = 99, p < 0.001, effect-size (r) = 0.83 |
| Fig. S6B | CS+/CS- vs CS3-/CS4-, odor period, Late session (vCA1) | Euclidean distance between MDS values | 10 MDS runs | Mann-Whitney U, bonferroni correction for n=2 | U = 0, p < 0.001, effect-size (r) = 0.85 |
| Fig. S6B | CS1+/CS2+ vs CS3-/CS4-, odor period, Late session (vCA1) | Euclidean distance between MDS values | 10 MDS runs | Mann-Whitney U, bonferroni correction for n=2 | U = 0, p < 0.001, effect-size (r) = 0.85 |
| Fig. S6B | CS+/CS- vs CS1+/CS2+, trace period, Late session (vCA1) | Euclidean distance between MDS values | 10 MDS runs | Mann-Whitney U, bonferroni correction for n=2 | U = 94, p = 0.002, effect-size (r) = 0.74 |
| Fig. S6B | CS+/CS- vs CS3-/CS4-, trace period, Late session (vCA1) | Euclidean distance between MDS values | 10 MDS runs | Mann-Whitney U, bonferroni correction for n=2 | U = 99, p < 0.001, effect-size (r) = 0.83 |
| Fig. S6B | CS1+/CS2+ vs CS3-/CS4-, trace period, Late session (vCA1) | Euclidean distance between MDS values | 10 MDS runs | Mann-Whitney U, bonferroni correction for n=2 | U = 75, p = 0.13, effect-size (r) = 0.42 |
| Fig. S6B | CS+/CS- vs CS1+/CS2+, US period, Late session (vCA1) | Euclidean distance between MDS values | 10 MDS runs | Mann-Whitney U, bonferroni correction for n=2 | U = 77, p = 0.09, effect-size (r) = 0.46 |
| Fig. S6B | CS+/CS- vs CS3-/CS4-, US period, Late session (vCA1) | Euclidean distance between MDS values | 10 MDS runs | Mann-Whitney U, bonferroni correction for n=2 | U = 0, p < 0.001, effect-size (r) = 0.85 |
| Fig. S6B | CS1+/CS2+ vs CS3-/CS4-, US period, Late session (vCA1) | Euclidean distance between MDS values | 10 MDS runs | Mann-Whitney U, bonferroni correction for n=2 | U = 93, p = 0.003, effect-size (r) = 0.73 |

| Figure | Variable | Unit of Comparison | n | Test | Results |
|---|---|---|---|---|---|
| Fig. S6C | CS1+/CS2+ vs CS3-/CS4-, US period, Late session (dCA1) | Euclidean distance between MDS values | 10 MDS runs | Mann-Whitney U, bonferroni correction for n=2 | U = 47, p = 1, effect-size (r) = 0.05 |
| Fig. S6C | CS+/CS- vs CS1+/CS2+, baseline period, Late session (dCA1) | Euclidean distance between MDS values | 10 MDS runs | Mann-Whitney U, bonferroni correction for n=2 | U = 58, p = 1, effect-size (r) = 0.14 |
| Fig. S6C | CS+/CS- vs CS3-/CS4-, baseline period, Late session (dCA1) | Euclidean distance between MDS values | 10 MDS runs | Mann-Whitney U, bonferroni correction for n=2 | U = 54, p = 1, effect-size (r) = 0.07 |
| Fig. S6C | CS1+/CS2+ vs CS3-/CS4-, baseline period, Late session (dCA1) | Euclidean distance between MDS values | 10 MDS runs | Mann-Whitney U, bonferroni correction for n=2 | U = 43, p = 1, effect-size (r) = 0.12 |
| Fig. S6C | CS+/CS- vs CS1+/CS2+, odor period, Late session (dCA1) | Euclidean distance between MDS values | 10 MDS runs | Mann-Whitney U, bonferroni correction for n=2 | U = 99, p < 0.001, effect-size (r) = 0.83 |
| Fig. S6C | CS+/CS- vs CS3-/CS4-, odor period, Late session (dCA1) | Euclidean distance between MDS values | 10 MDS runs | Mann-Whitney U, bonferroni correction for n=2 | U = 77, p = 0.09, effect-size (r) = 0.46 |
| Fig. S6C | CS1+/CS2+ vs CS3-/CS4-, odor period, Late session (dCA1) | Euclidean distance between MDS values | 10 MDS runs | Mann-Whitney U, bonferroni correction for n=2 | U = 13, p = 0.012, effect-size (r) = 0.63 |
| Fig. S6C | CS+/CS- vs CS1+/CS2+, trace period, Late session (dCA1) | Euclidean distance between MDS values | 10 MDS runs | Mann-Whitney U, bonferroni correction for n=2 | U = 100, p < 0.001, effect-size (r) = 0.85 |
| Fig. S6C | CS+/CS- vs CS3-/CS4-, trace period, Late session (dCA1) | Euclidean distance between MDS values | 10 MDS runs | Mann-Whitney U, bonferroni correction for n=2 | U = 100, p < 0.001, effect-size (r) = 0.85 |
| Fig. S6C | CS1+/CS2+ vs CS3-/CS4-, trace period, Late session (dCA1) | Euclidean distance between MDS values | 10 MDS runs | Mann-Whitney U, bonferroni correction for n=2 | U = 89, p = 0.007, effect-size (r) = 0.66 |
| Fig. S6C | CS+/CS- vs CS1+/CS2+, US period, Late session (dCA1) | Euclidean distance between MDS values | 10 MDS runs | Mann-Whitney U, bonferroni correction for n=2 | U = 100, p < 0.001, effect-size (r) = 0.85 |
| Fig. S6C | CS+/CS- vs CS3-/CS4-, US period, Late session (dCA1) | Euclidean distance between MDS values | 10 MDS runs | Mann-Whitney U, bonferroni correction for n=2 | U = 100, p < 0.001, effect-size (r) = 0.85 |
| Fig. S6C | CS1+/CS2+ vs CS3-/CS4-, US period, Late session (dCA1) | Euclidean distance between MDS values | 10 MDS runs | Mann-Whitney U, bonferroni correction for n=2 | U = 47, p = 1, effect-size (r) = 0.05 |
| Fig. S7B | CS+Shock/CS- decoding accuracy, odor period, Early vs Late (vCA1) | 10 decoding iterations for each session | n-matched pseudopopulation of 444 cells from 10 vCA1 mice | Mann-Whitney U | U = 0, p < 0.001, effect-size (r) = 0.85 |
| Fig. S7B | CS+Shock/CS- decoding accuracy, odor period, Early vs Late (dCA1) | 10 decoding iterations for each session | n-matched pseudopopulation of 444 cells from 3 dCA1 mice | Mann-Whitney U | U = 49.5, p = 1, effect-size (r) = 0.008 |
| Fig. S7B | CS+Shock/CS- decoding accuracy, trace period, Early vs Late (vCA1) | 10 decoding iterations for each session | n-matched pseudopopulation of 444 cells from 10 vCA1 mice | Mann-Whitney U | U = 9, p = 0.002, effect-size (r) = 0.69 |
| Fig. S7B | CS+Shock/CS- decoding accuracy, trace period, Early vs Late (dCA1) | 10 decoding iterations for each session | n-matched pseudopopulation of 444 cells from 3 dCA1 mice | Mann-Whitney U | U = 12, p = 0.005, effect-size (r) = 0.64 |
| Fig. S7C | CS+Rew/CS- decoding accuracy, odor period, Early vs Late (vCA1) | 10 decoding iterations for each session | n-matched pseudopopulation of 444 cells from 10 vCA1 mice | Mann-Whitney U | U = 0, p < 0.001, effect-size (r) = 0.85 |
| Fig. S7C | CS+Rew/CS- decoding accuracy, odor period, Early vs Late (dCA1) | 10 decoding iterations for each session | n-matched pseudopopulation of 444 cells from 3 dCA1 mice | Mann-Whitney U | U = 0, p < 0.001, effect-size (r) = 0.77 |
| Fig. S7C | CS+Rew/CS- decoding accuracy, trace period, Early vs Late (vCA1) | 10 decoding iterations for each session | n-matched pseudopopulation of 444 cells from 10 vCA1 mice | Mann-Whitney U | U = 11, p = 0.004, effect-size (r) = 0.66 |
| Fig. S7C | CS+Rew/CS- decoding accuracy, trace period, Early vs Late (dCA1) | 10 decoding iterations for each session | n-matched pseudopopulation of 444 cells from 3 dCA1 mice | Mann-Whitney U | U = 0, p < 0.001, effect-size (r) = 0.85 |
| Fig. S7G | Odor identity decoding accuracy across reversal learning (Late/Late Reversal), vCA1 vs dCA1 | 10 decoding iterations for each region | n-matched pseudopopulation of 281 cells from 10 vCA1 and 3 dCA1 mice | Mann-Whitney U | U = 41, p = 0.52, effect-size (r) = 0.15 |
| Fig. S7H | Trial type decoding accuracy across reversal learning, trace period, Rew/CS- accuracy vs Sh/CS- accuracy (vCA1) | 10 decoding iterations each | n-matched pseudopopulation of 281 cells from 10 vCA1 mice | Mann-Whitney U | U = 100, p < 0.001, effect-size (r) = 0.85 |
| Fig. S7H | Trial type decoding accuracy across reversal learning, trace period, Rew/CS- accuracy vs Sh/CS- accuracy (dCA1) | 10 decoding iterations each | n-matched pseudopopulation of 281 cells from 3 dCA1 mice | Mann-Whitney U | U = 100, p < 0.001, effect-size (r) = 0.85 |
